# Engineering a cell-free biosensor signal amplification circuit with polymerase strand recycling

**DOI:** 10.1101/2024.04.25.591074

**Authors:** Yueyi Li, Tyler Lucci, Matias Villarruel Dujovne, Jaeyoung Kirsten Jung, Daiana A. Capdevila, Julius B. Lucks

## Abstract

Cell-free systems are powerful synthetic biology technologies because of their ability to recapitulate sensing and gene expression without the complications of living cells. Cell-free systems can perform even more advanced functions when genetic circuits are incorporated as information processing components. Here we expand cell-free biosensing by engineering a highly specific isothermal signal amplification circuit called polymerase strand recycling (PSR) that leverages T7 RNA polymerase off-target transcription to recycle nucleic acid inputs within DNA strand displacement circuits. We develop design rules for PSR circuit components and use these rules to construct modular biosensors that can directly sense different RNA targets with limits of detection in the nM range and high specificity. We then use PSR for signal amplification within allosteric transcription factor-based biosensors for small molecule detection. We use a double equilibrium model of transcription factor:DNA and transcription factor:ligand binding interactions to predict biosensor sensitivity enhancement by PSR, and then demonstrate this approach experimentally by achieving 3.6-4.6-fold decreases in biosensor EC50 to sub micromolar ranges. We believe this work expands the current capabilities of cell-free circuits by incorporating PSR, which we anticipate will have a wide range of uses within biotechnology.

## Introduction

Cell-free systems have enabled rapid, point-of-care detection of a wide range of important environmental and human health targets such as metals [1–3] (arsenic, lead, copper, cadmium, zinc, mercury), ions and small molecules [4–6] (fluoride, vitamin B12, benzoic acid derivatives), antibiotics [2] (tetracycline, oxytetracycline, chlortetracycline), disinfectant bi- products [2] (benzalkonium chloride), viruses [7] (Zika, Ebola), agricultural toxins [8] (atrazine), drugs [9, 10] (cocaine, gamma-hydroxybutyrate) and microbial quorum sensing molecules [11]. Furthermore, cell-free biosensing reactions can be assembled, freeze-dried, and simply rehydrated with samples for usage [12–15], and can be incorporated into device formats that can be used by non-experts [16]. As such, these systems are safe, easy-to-use, and field-deployable, showing great promise to address societal needs in environmental monitoring and human health [2, 5, 7, 9].

While promising, it is still a challenge to engineer cell-free biosensors to be able to detect chemicals at low enough concentrations to meet practical or regulatory use requirements, with recent reports of allosteric transcription factor (aTF)-based cell-free biosensors demonstrating limits of detection in the µM to mM range [2, 17, 18], while many toxic chemicals are regulated at the nM level [2, 17–19]. To date, many strategies have been pursued for improving cell-free biosensor sensitivity such as enzymatic amplification or interfacing biosensors with physical devices such as electrochemical devices or lateral flow assays [20].

Electrochemical biosensing has the potential to increase sensitivity [21–23], but assay development is complex and expensive and often requires further miniaturization [20]. Lateral flow assays can improve limit of detection by decreasing the sample flow rate and increasing the duration of affinity reactions [24–26], but can be prone to interference by other analytes [20]. Enzymatic amplification systems such as isothermal nucleic acid amplification combined with CRISPR-Cas detection have been adapted for rapid sensing of nucleic acids and have reached the aM range for sensitivity [27]. However, these systems require extensive sample preparation steps [27], and are not immediately applicable to the detection of small molecule compounds.

Here we sought to develop a simpler approach to improving cell-free biosensor sensitivity that leverages our ability to program nucleic acid circuits. Cell-free biosensors typically consist of an aTF that is designed to bind to a DNA reporter template: binding of the aTF to the DNA blocks expression of the reporter molecule, while the presence of a target ligand that binds to the aTF removes this block and allows reporter expression [2]. Previously, we showed that biosensors could be made more sensitive by simply reducing the amount of aTF in these reactions [2]. However, this strategy resulted in a significant increase in background signal (i.e., signal in the absence of ligand) due to insufficient repression of the DNA reporter template [2]. We reasoned that this could be overcome by introducing a signal amplification layer to the system that would allow lower amounts of DNA template to be used, resulting in lower background in the absence of ligand and strong signals when small amounts of ligand are present.

Our recent development of a cell-free small molecule sensing platform called RNA Output Sensors Activated by Ligand INDuction (ROSALIND) [2] that utilizes toehold mediated strand displacement (TMSD) circuits [28] creates the possibility for such a signal amplification layer by leveraging the natural ability of T7 RNA polymerase to transcribe from 3’ toeholds exposed on DNA duplexes [28, 29]. Specifically, we developed a new concept in nucleic acid circuitry, called polymerase strand recycling (PSR), that uses T7 RNAP off-target transcription from 3’ toeholds to recycle TMSD circuit inputs during the course of a biosensing reaction.

ROSALIND interfaced with PSR consists of three core layers: a sensing layer, a processing layer, and signal amplification layer (Figure 1). The sensing layer generates an RNA strand as a result of target ligand sensing. The processing layer takes the sensing layer RNA strand, utilizes TMSD circuits to add computational features if desired [28], and generates an output DNA strand. The signal amplification layer consists of a quenched fluorophore DNA TMSD gate that can be strand displaced by the output of the processing layer. This is designed to result in a 3’ overhanging toehold which can be transcribed by T7 RNAP off-target transcription, resulting in the release of the original DNA strand, thus recycling it for further rounds of signal generation to enable signal amplification.

**Figure 1.**
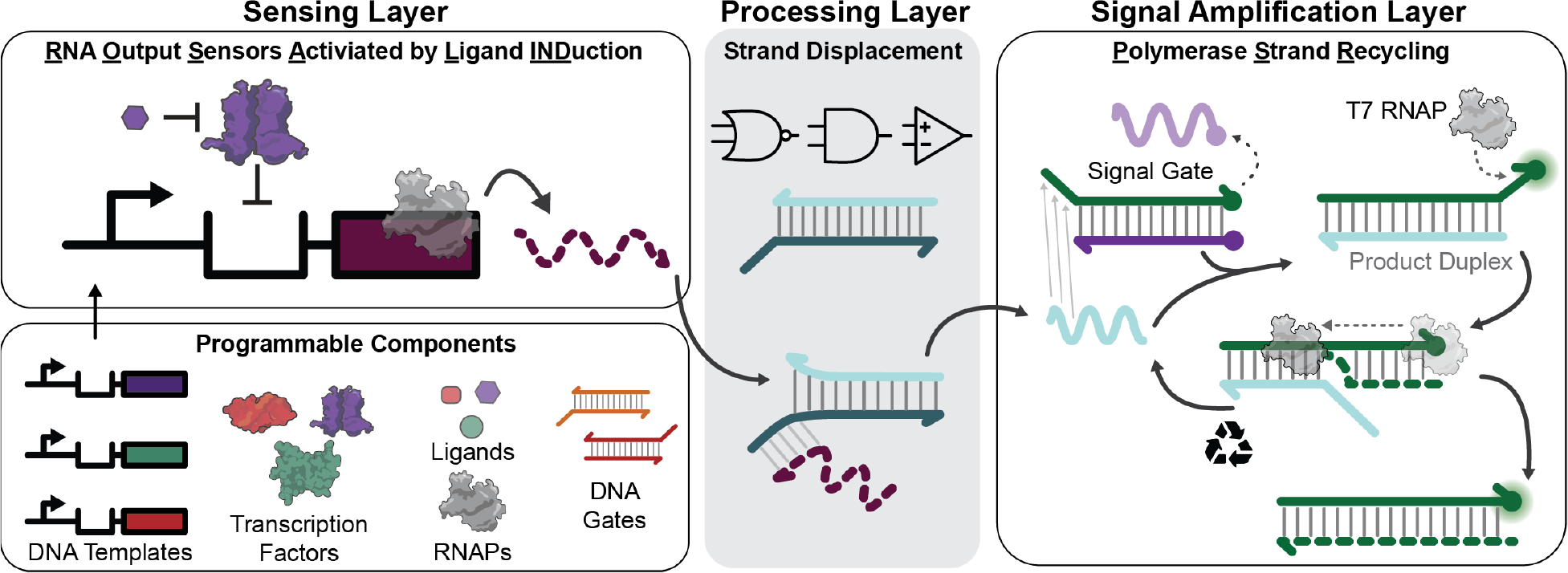
Amplifying outputs of programmable cell-free biosensor circuits with polymerase strand recycling. (Left) Programmable cell-free biosensor circuits are configured by genetically wiring multiple components together, including transcription factors for sensing, regulated DNA templates for outputs, RNA polymerases (RNAPs) to drive circuit function, and chemical ligands for detection. Biosensing circuits activate when a ligand binds to a protein transcription factor, causing it to un-bind from a DNA template and activate transcription of RNA output signals. (Middle) RNA output signals can then interact with programmable DNA gates via toehold-mediated strand displacement (TMSD). DNA gates can be configured to process biosensor signals by adding computational features such as logic processing and signal comparison. (Right) Outputs of the processor layer generate a detectable signal by strand displacing a quenched fluorophore signal gate. These output signals can be further amplified with polymerase strand recycling (PSR), a process by which RNAP transcribes the resulting bound fluorophore strand, thus releasing a strand that can invade a new signal gate to create a cycle of signal amplification.

Here, we develop PSR by exploring the design principles of this architecture and applying it to sensitize circuits for the detection of both nucleic acid and small molecule ligands. We start by investigating the PSR phenomenon using RNA and DNA inputs, and show that T7 RNAP-mediated strand recycling only occurs when DNA TMSD gates are used. We next show that PSR can be used to amplify the signals from transcriptional circuits by designing an intermediate fuel gate as part of the processing layer to convert an input RNA into a recyclable DNA molecule that can be regenerated through PSR. We show this circuit design can be directly applied to the detection of RNA inputs, and demonstrate a limit of detection (LoD) of 5 - 250 nM for model miRNA targets. Interfacing PSR with ROSALIND by incorporating aTF operator sequences in the transcriptional templates allows PSR circuits to amplify signals from biosensing circuits that detect small molecules. We show that the incorporation of PSR allows a reduction in both the aTF and DNA template concentration required to enact sensing function, while still producing a signal compatible with on-site readout. Using a double equilibrium model of aTF:DNA and aTF:ligand binding interactions, we predict that lowering aTF concentration can potentially improve sensitivity characterized by EC50 by up to three orders of magnitude if DNA template concentration is also sufficiently lowered to ensure adequate repression. We experimentally demonstrate how PSR enables this approach by applying it to the tetracycline sensing system using the TetR aTF to achieve an EC50 of 0.07 µM, representing a 3.6-fold reduction in EC50 from TMSD sensor circuits that do not use PSR. We also apply this approach to the zinc sensing system using the SmtB aTF to achieve an EC50 of 0.46 µM, representing a 4.6-fold reduction in EC50 from TMSD sensor circuits that do not use PSR. Importantly, these results represent a respective 14.7- and 15.1-fold reduction in EC50 over previously reported values for tetracycline and zinc detection [28]. Overall, this work shows that PSR is a generalizable method that can amplify cell-free biosensor signals and be used to improve the sensitivity of detection of a range of target ligands.

## Results

### Establishing Polymerase Strand Recycling with T7 RNA Polymerase

We first sought to test the principle of PSR by designing an amplification system using T7 RNAP. As T7 RNAP is well characterized to transcribe DNA duplexes that contain 3’ toeholds without its natural promoter sequence [28–30], it should be possible to generate a signal amplification system within TMSD circuits by leaving 3’ toeholds on the product duplexes of strand displacement reactions (Figure 1).

TMSD signal generation systems typically consist of two components: an output nucleic acid strand produced directly from the sensing layer, and a fluorophore-strand:quencher-strand DNA signal gate (Figure 1). Strand displacement of the signal gate by the output strand results in a fluorophore-strand:output duplex that can generate fluorescent signal. To test the concept of PSR, we designed two ‘recycle’ strands to strand displace a DNA signal gate from previous work [31] and leave a four-nucleotide (nt) toehold on the 3’-end of the fluorophore strand. Both RNA (RNA input) and DNA (DNA input) versions of the recycle strands were designed to investigate potential differences in the ability of T7 RNAP to transcribe RNA-DNA and DNA-DNA duplexes from 3’ overhangs (Figure 2a).

**Figure 2.**
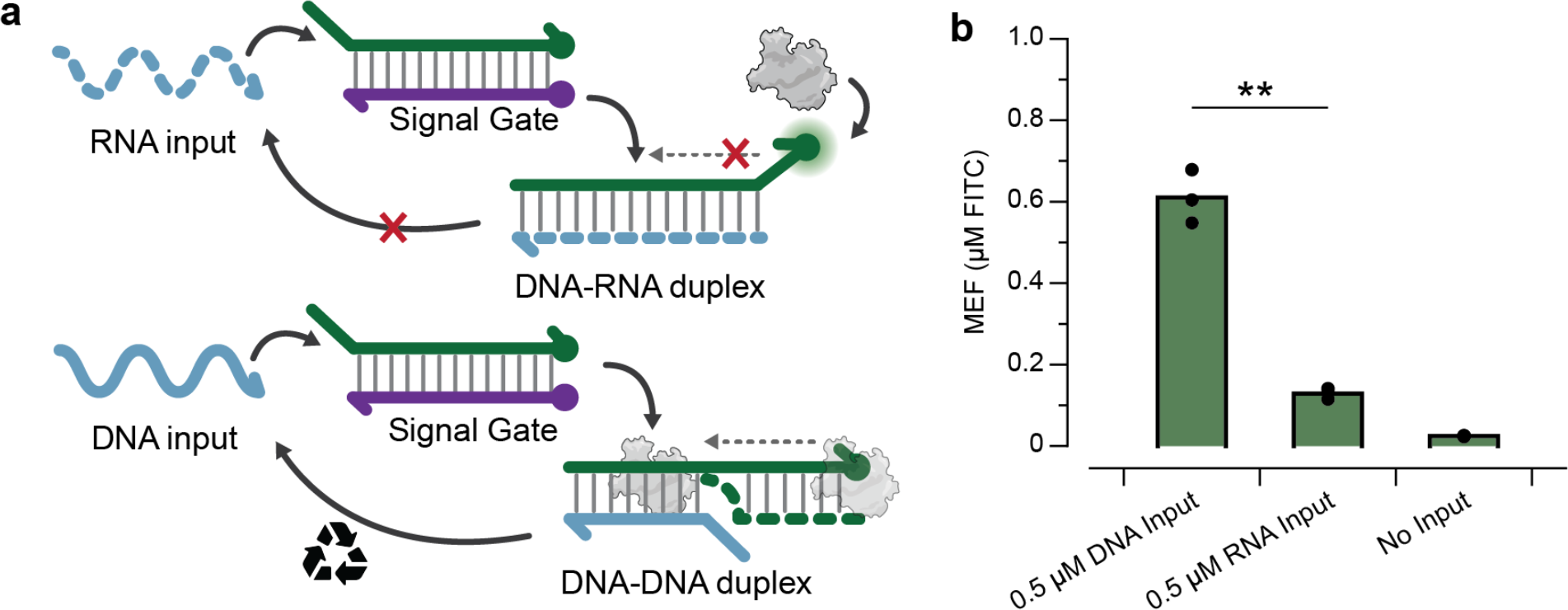
A DNA-DNA duplex is required for T7 RNAP off-target transcription. a. Invading RNA (RNA input) or DNA (DNA input) strands were designed to strand displace the DNA signal gate and form a new duplex with the fluorophore-strand, leaving a 4-nt 3’ toehold. Transcription of the resulting 3’ toehold by T7 RNAP should result in signal amplification. **b**. Fluorescent read outs of end-point reactions (1 hr) consisting of 2.5 µM of the signal gate, 2 ng T7 RNAP, and either 0.5 µM of RecycleD, RecycleR or no input. RecycleD is able to achieve significantly higher signal than RecycleR. Data shown are n = 3 experimentally independent replicates, each plotted as a point with raw fluorescence standardized to MEF (µM FITC). Bar heights represent the average over these replicates. A paired Student’s t-test was used to compare the fluorescent signals between RecycleR and RecycleD. The p-value range is indicated by asterisks (***p < 0.001, **p = 0.001–0.01, *p = 0.01–0.05). Exact p-value can be found in Source Data.

DNA and RNA inputs were purified and added to IVT reactions containing 2.5 µM of the DNA signal gate, 2 ng T7 RNAP, and 0.5 µM of either DNA input, RNA input, or no input (Figure 2b). In the presence of RNA input, we observed low fluorescence activation (Figure 2b), while with DNA input, the fluorescence activation reached a level that was close to that of 2.5 µM of isolated fluorophore-strand (Figure 2b, Supplementary Figure 2b). This suggests that RNA input can strand displace the signal gate, but T7 RNAP cannot recycle it efficiently from the resulting DNA-RNA hybrid fluorophore-strand:RNA-input complex. In contrast, the high signal observed from using DNA input suggests that T7 RNAP can transcribe the DNA-DNA fluorophore- strand:DNA-input complex.

Overall, these results show that PSR is possible, though only with DNA-DNA duplexes, which informs the implementation of PSR within biosensing circuits.

### Adding a “fuel gate” to enable signal amplification through polymerase strand recycling

The above results showed that PSR only works using a DNA strand as the recyclable output signal. However, transcriptional biosensors only produce RNA outputs, necessitating an interface that can convert an RNA signal into a DNA strand. To create this interface, we designed a “fuel gate” [32] that consisted of a DNA duplex comprised of a DNA strand, RecycleD and a complementary strand, designed to exchange a generated RNA strand, RecycleR, with RecycleD via toehold-mediated strand displacement (Figure 3a, Supplementary Figure 3). In this way, transcription of the RecycleR sequence can in principle invade and strand displace the fuel gate, releasing RecycleD for participation in downstream PSR amplification.

**Figure 3:**
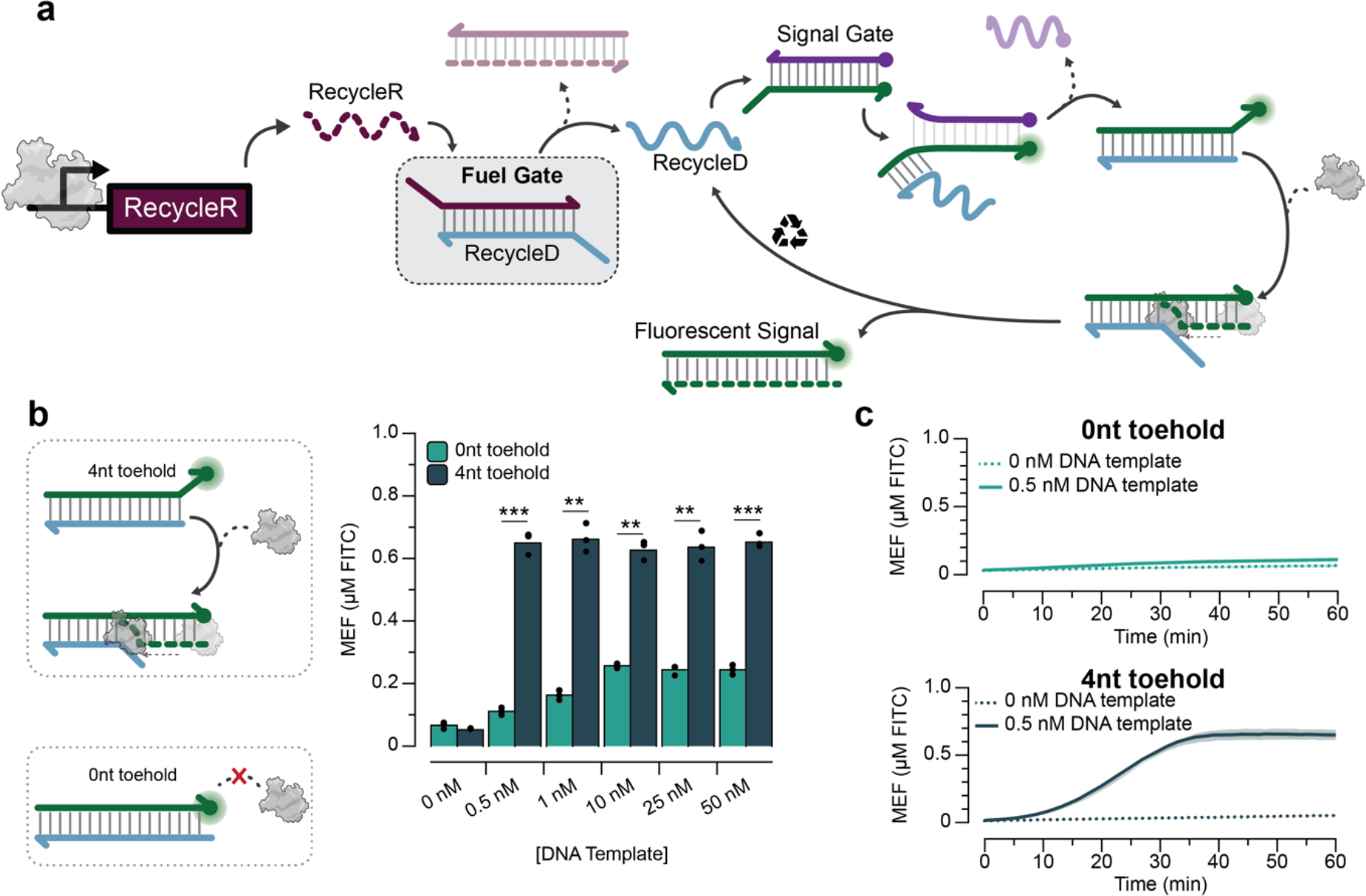
Interfacing PSR amplification with transcriptional signals via a “fuel gate”. a. The fuel gate concept. The fuel gate is designed to sequester RecycleD with a complementary strand, leaving 5’ toehold overhangs. Transcription of RecycleR can then lead to invasion of the fuel gate, releasing RecycleD for PSR. **b.** Toehold length for PSR affects signal amplification. With a designed 4-nt 3’ toehold on the RecycleD:fluorophore duplex, greater fluorescence activation was observed after 1 hr from a circuit containing 1.5 µM of fuel gate, 2.5 µM of signal gate, 2 ng T7 RNAP and 0.5 nM – 50 nM of DNA transcription template encoding RecycleR, compared to the same circuit with a 0-nt 3’ toehold. In addition, the 4-nt toehold led to a plateau of signal at 0.5 nM transcription template compared to 10 nM for the 0-nt toehold. The DNA template concentrations at which the 4-nt signals are distinguishable from 0-nt signals were determined using paired Student’s t-tests. The p-value range is indicated by asterisks (***p < 0.001, **p = 0.001–0.01, *p = 0.01–0.05). Exact p-values can be found in Source Data. **c.** Kinetic data for 0 nM and 0.5 nM DNA template concentrations with and without a 4nt toehold. Raw fluorescence data are standardized to MEF (µM FITC). Data shown in b are n = 3 experimentally independent replicates, each plotted as a point. Bar heights represent the average over these replicates. Data shown in c are n = 3 experimentally independent replicates. The lines and dotted lines represent the average of the replicates. Shading indicates the average of the replicates ± s.d.

To test this concept, we designed DNA templates encoding the T7 RNAP promoter followed by two initiating guanines and the RecycleR sequence (Supplementary Data).

Reactions were then assembled containing 2 ng T7 RNAP, 1.5 µM of the fuel gate, 2.5 µM of the signal gate and either 0 nM or 1 nM of the DNA template (Supplementary Figure 2).

Interestingly, we observed that T7 RNAP could activate fluorescence signal even without the presence of DNA template (Supplementary Figure 2). We also observed that the signal activation did not reach the full level of the fluorescence observed only with 2.5 µM of the fluorophore strand alone, which we attribute to TMSD being unable to fully release 2.5 µM of the fluorophore strand from the quencher strand, as well as freed quencher strands still sequestering some fluorescent signal.

We hypothesized that the signal observed even without DNA transcription template was caused by the fuel gate duplex being too weak, allowing a small amount of dissociated RecycleD strand to displaced DNA signal gates and initiate PSR. To mitigate this effect, we incorporated 2’-O-methyl modifications on both fuel and signal gates, which are known to stabilize nucleic acid duplexes [33] (Supplementary Figure 4). Testing different combinations of modified fuel and signal gates showed that incorporating 2’-O-methyl modified fuel gates significantly reduced the background signal in the absence of DNA template (Supplementary Figure 4).

Having demonstrated a functioning system, we next investigated how the system’s response varied with DNA template concentration, as well as the importance of the 4nt toehold of the signal gate. Reactions were set up as before, but containing a variable amount of DNA template in the range of 0.5-50 nM, and using versions of RecycleD designed to leave either 4- nt or 0-nt 3’ toeholds on the fluorophore-strand:RecycleD complex (Figure 3b). As expected, we observed significantly higher fluorescent signal with the 4-nt toehold than the 0-nt toehold (Figure 3b). In addition, for the 4-nt toehold case, we observed full signal generation with as little as 0.5 nM DNA template (Figure 3b), which is markedly lower than the 10-25 nM used in previous versions of the ROSALIND system [2, 28]. An examination of reaction kinetics showed that this signal amplification can be achieved in ∼35 minutes (Figure 3c), similar to the signal generation times in previous versions of ROSALIND [28].

Overall, these results demonstrate that PSR can process RNA inputs and serve as an amplification system for transcriptional circuits through the inclusion of a fuel gate, and that doing so allows a significant reduction in DNA template while preserving reaction kinetics with similar fluorescent output.

### PSR circuits can be used to directly sense RNA targets

After demonstrating that PSR can generate a fluorescent signal from RNA molecules transcribed from DNA templates, we next sought to explore the detection of RNA targets that are directly input into the system. Specifically, we modified the sequences of the fuel and signal gates to match portions of desired target RNAs where the target RNAs can release RecycleD for downstream PSR amplification (Supplementary Figure 5). The fuel gate contains a 2’-O-methyl modified strand that is complementary to the RNA target, plus several additional nucleotides to form a ∼20-bp overlap with the unmodified RecycleD strand. The signal gate requires a fluorophore-strand that contains complementary sequence to RecycleD but leaves a 4-nt 3’ toehold when displaced to allow PSR, and a quencher-strand that is complementary to the fluorophore-strand but leaves an 8nt 5’ toehold on the fluorophore-strand to allow efficient invasion by RecycleD (Figure 4a). In this way, inputting the desired RNA signal should strand invade the fuel gate and release RecycleD to generate and amplify a fluorescent signal.

**Figure 4.**
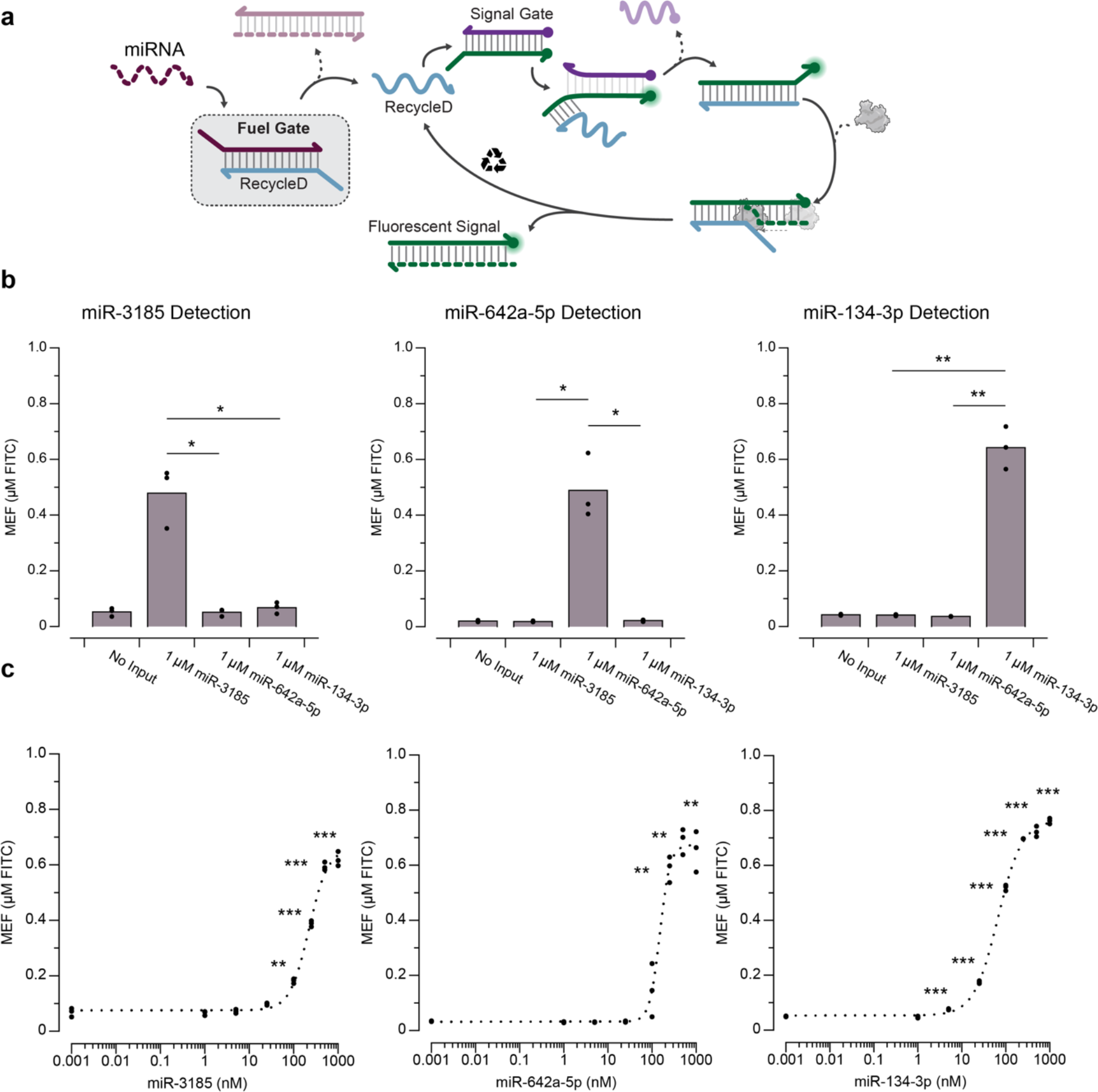
Nucleic acid sensing with PSR. a. PSR for detecting miRNAs. Fuel gates are designed to interact directly with miRNA targets, releasing RecycleD for PSR. **b.** PSR miRNA sensing specificity. Different sequences for PSR fuel gates and signal gates were designed to sense miRNA3185, miRNA642a-5p, and miRNA134-3p. The PSR circuits containing 1.5 µM of fuel gate, 2.5 µM of signal gate, 2 ng T7 RNAP all detect 1 µM of their respective miRNA targets with significant specificity after 2 hrs. Significance values comparing fluorescence from targeted miRNA inputs and the indicated non-targeted inputs were determined using a paired Student’s t-test. The p-value range is indicated by asterisks (***p < 0.001, **p = 0.001–0.01, *p = 0.01–0.05). Exact p-values can be found in Source Data. **c.** PSR miRNA sensing limit of detection. Dose response with miRNA3185 measured after 2 hrs shows limit of detection (LoD) of 100 nM. Dose response with miRNA642a-5p measured after 2 hrs shows LoD of 250 nM. Dose response with miRNA3185 measured after 2 hrs shows LoD of 5 nM. The LoDs were determined from the concentration value at which the signal was significantly greater than the no input condition using a paired Student’s t-test. The p-value range is indicated by asterisks (***p < 0.001, **p = 0.001–0.01, *p = 0.01–0.05). Exact p-values can be found in Source Data. Data shown in b are n = 3 experimentally independent replicates, each plotted as a point, with bar heights representing the average over these replicates. Data shown in c are n = 3 experimentally independent replicates, each plotted as a point with raw fluorescence standardized to MEF (µM FITC). Dotted curves in c represent Hill Equation fits (see Methods), with fit parameters in Source Data.

As a model target, we chose microRNAs (miRNAs), short non-coding RNAs that play an important role in gene regulation and have been widely reported as potential biomarkers for human diseases [34, 35], thus making them valuable targets for biosensors [36–38]. Specifically, we chose three distinct miRNAs (miR-3185, miR-642a-5p, and miR-134-3p) that are biomarkers for fatigue and mood disturbances as inputs for PSR [39].

As a proof-of-concept, miRNA targets were synthesized and purified, and specificity of the platform was tested by adding 1 µM of input to each of three PSR circuits configured to sense only a single miRNA. Fluorescence characterization demonstrated that all three PSR systems were able to generate significantly higher fluorescent signal with their respective miRNA compared to the other two miRNAs (Figure 4b).

We next assessed sensitivity of the system by performing a titration with miRNAs and found the LoD, defined as the concentration value at which the signal was significantly greater than the no input condition, to be within the 5 – 250 nM range for PSR to detect nucleic acid targets (Figure 4c). Specifically, the miR-3185 target was detected with a LoD of 100 nM, the miR-642a-5p with a LoD of 250 nM, and the miR-134-3p with a LoD of 5 nM. Differences may be attributed to the number of base pairs in the fuel gate design (Supplementary Figure 5).

Overall, these results demonstrate that PSR circuits can be used to directly sense RNA inputs with high specificity and LoDs between 5 - 250 nM.

### Interfacing transcriptional biosensors with PSR signal amplification

We next sought to interface the PSR platform with transcriptional regulation by allosteric transcription factors (aTFs). With appropriately configured reporter templates, aTFs can bind to their cognate operator sequences and repress downstream transcription by T7 RNAP in the absence of its cognate ligand. However, in the presence of an aTF’s cognate ligand, the aTF unbinds from its operator sequence, allowing downstream transcription. We previously showed that this setup can be used as a cell-free biosensing platform by swapping in different aTF/operator pairs to detect different chemical contaminants in water [28]. In addition, we showed that this platform can be coupled to strand displacement circuitry by configuring the transcribed output RNA molecules to be complementary to downstream DNA gates, creating an opportunity to interface aTF biosensors with PSR circuitry.

To explore whether PSR signal amplification can be regulated with an aTF, we chose to focus on the TetR repressor by inserting the *tetO* operator sequence in between the T7 RNAP promoter and RecycleR, using a 2-bp spacer between the two sequences that was found in previous work to be optimal for TMSD signal generation [28] (Figure 5a). We first tested whether the addition of *tetO* impacted PSR signal generation without TetR. Titrating a range of input DNA template concentrations revealed full signal generation with as little as 0.5 nM DNA template (Figure 5b), representing a 20X reduction compared to the 10 nM DNA template needed to generate signal from an equivalent TMSD reaction without PSR circuitry (Supplementary Figure 6).

**Figure 5.**
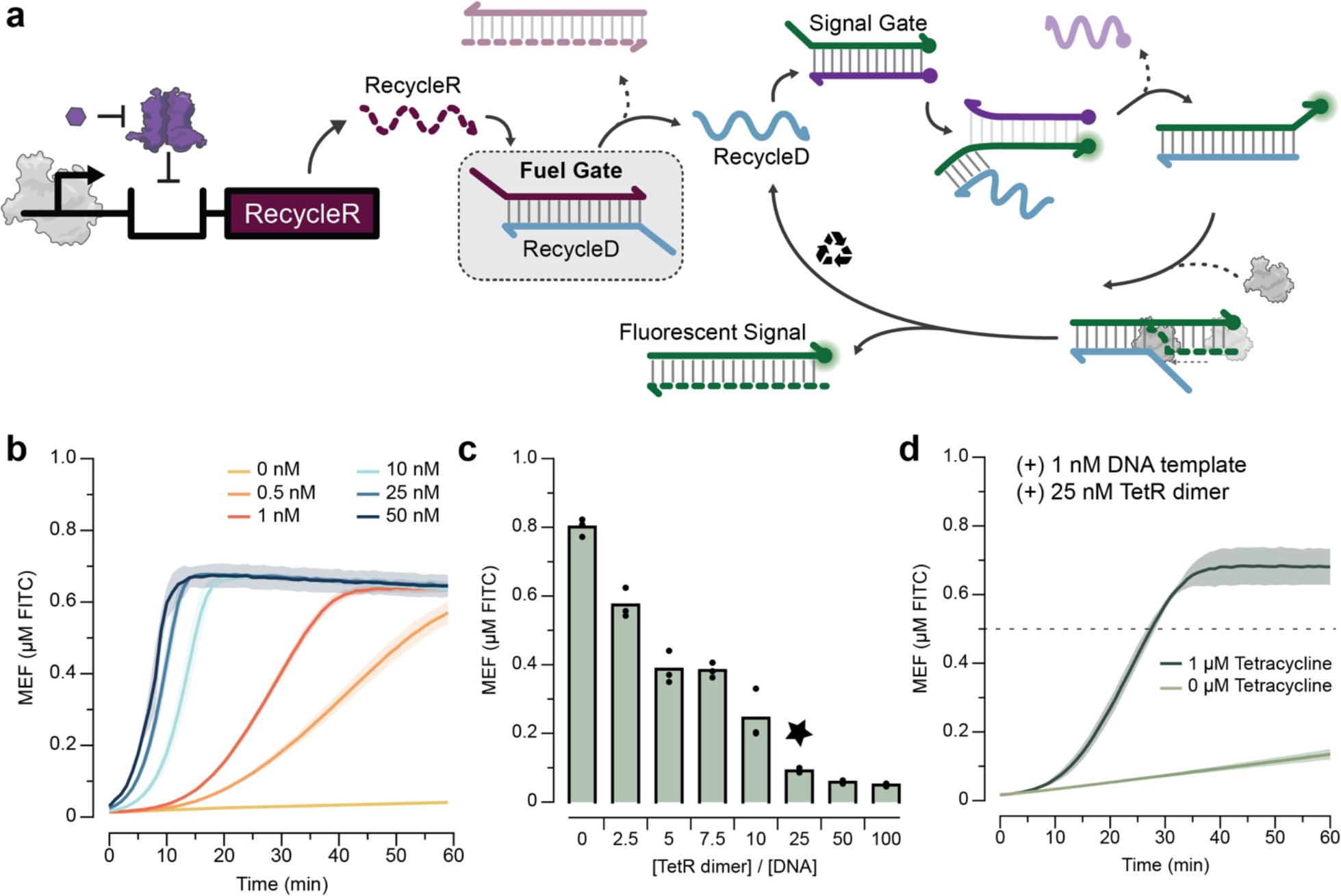
Interfacing transcriptional biosensors with PSR signal amplification. a. RecycleR transcription can be allosterically regulated with a DNA template configured to bind a purified aTF (here TetR) via an operator sequence (here *tetO*) placed downstream of the T7 promoter and upstream of the RecycleR coding sequence. Activation of the aTF biosensor in the presence of a target ligand (here aTc) generates a signal which is amplified by the downstream PSR circuit. **b.** Kinetic traces of the circuit containing 1.5 µM of fuel gate, 2.5 µM of signal gate, 2 ng T7 RNAP and 0 nM – 50 nM of RecycleR DNA transcription templates containing the *tetO* sequence, showing that even the lowest template concentration (0.5 nM) can reach nearly maximum fluorescence activation within 1 hr **c.** Fluorescence levels after 1 hr for circuits containing 0.5 nM of DNA transcription template and a range of input TetR concentrations from 0 – 100 nM. A ratio of TetR dimer:DNA template of 25:1 (12.5 nM:0.5 nM, starred) was found to be the minimal amount of TetR needed to completely repress 0.5 nM of DNA template. **d.** Kinetic traces of the complete biosensor-PSR circuit containing 1 nM of DNA transcription template and 25 nM of TetR dimer, with and without 1 µM of added tetracycline. The dashed line at 0.5 µM FITC represents the visible fluorescence threshold found previously [28]. Raw fluorescence data are standardized to MEF (µM FITC). Data shown in b and d are n = 3 experimentally independent replicates for each condition, with lines representing the average of these replicates. Shading indicates the average of the replicates ± s.d. Data shown in c are n = 3 experimentally independent replicates, each plotted as a point, with bar heights representing the average over these replicates.

We next determined the amount of TetR dimer required to fully repress the PSR and TMSD-only systems. For purposes of comparing regulated PSR vs. TMSD reactions, we chose to use 0.5 nM of PSR DNA template and 10 nM of TMSD DNA template which generated full reaction signals (Supplementary Fig 6). Titrating in a range of TetR concentrations showed that 0.5 nM of PSR DNA template can be fully repressed by 12.5 nM TetR dimer (25X) (Figure 5c), while 50 nM of TetR dimer (5X) was required to repress the TMSD circuit (Supplementary Figure 7).

Finally, we tested whether the PSR signal amplification circuitry could be de-repressed by introducing the cognate ligand, tetracycline, to the TetR-regulated PSR system. Kinetic reaction traces demonstrate that the addition of tetracycline can induce transcription of RecycleR and activate fluorescent signal in a PSR reaction in a similar amount of time (∼30 min) as previously observed in regulated TMSD reactions (Figure 5d) [28].

Overall, these results demonstrate that PSR can be coupled with aTF regulation to yield a functional biosensor signal amplification system. Intriguingly, the use of PSR also allows the reduction of the amount of DNA template and aTF needed to activate full fluorescent signal in the presence of ligand.

### A simple equilibrium model of aTF:Ligand and aTF:DNA interactions predicts improvements in biosensor sensitivity

The previous results demonstrate that PSR amplified biosensor circuits reduce the concentrations of aTF and DNA template required to enact sensing function. Previous work showed that reducing the concentration of aTF in the system can sensitize cell-free biosensors, though this strategy had the undesirable effect of increasing background signal in the absence of ligand because DNA template concentration was not sufficiently reduced, resulting in insufficient repression [2]. Because PSR reduces both the required DNA template concentration and aTF concentration to enact sensing function, we hypothesized that PSR signal amplification circuitry could achieve the effect of increased sensitivity as observed in previous work without suffering from increased background signal in the absence of ligand.

To investigate this hypothesis, we constructed a simple double-equilibrium model describing the behavior of aTF:ligand and aTF:DNA binding interactions (see Supplementary Note). The model assumes that the timescale of transcription is much longer than the timescales of aTF:Ligand and aTF:DNA binding, which exist at equilibrium. The model takes as input DNA template concentration, aTF concentration, aTF:ligand dissociation constant, aTF:DNA dissociation constant, and a range of ligand concentrations. The model then solves for the fraction of DNA template without aTF bound, which we assume directly relates to the level of transcription output.

To study the tetracycline-TetR system, we used literature values for the TetR:tetracycline and TetR:DNA dissociation constants. We used the model to generate dose response curves for a range of TetR and DNA template concentrations. The model predicts that decreasing TetR concentration at constant TetR/DNA ratio can improve sensitivity characterized by EC50, the amount of ligand required to produce half-maximal induction, by approximately three orders of magnitude (Figure 6a). However, when TetR concentration approaches the TetR-DNA dissociation constant (∼0.014 nM), predicted leak begins to increase where a non-negligible fraction of DNA template remains free from TetR repression in the absence of appreciable ligand (Figure 6a). The TetR:DNA dissociation constant thus limits the extent to which lowering TetR concentration can be used to reduce EC50 without resulting in increased background signal. Accordingly, the model predicts that lowering the TetR:DNA dissociation constant can allow further reductions in EC50 without increasing background signal, as long as TetR will still preferentially bind ligand over DNA, or in other words, as long as the TetR:tetracycline dissociation constant is sufficiently lower than the TetR:DNA dissociation constant (Figure 6b). Overall, these results suggest that the ability of PSR-amplified reactions to reduce aTF concentration should lead to improvements in biosensor sensitivity.

**Figure 6.**
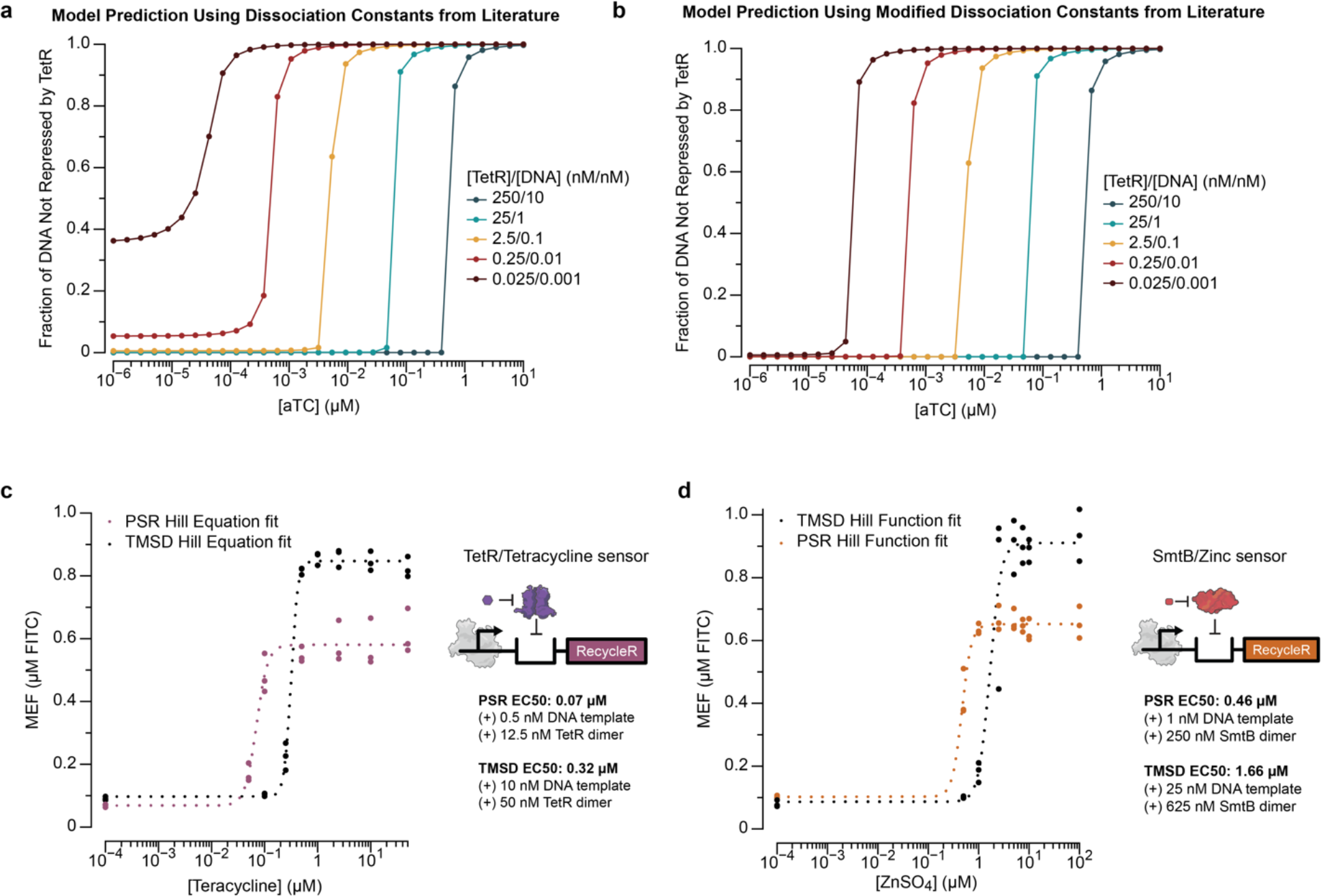
Polymerase strand recycling circuits can sensitize cell-free biosensors. a. The double equilibrium model predicts that decreasing aTF concentration at a constant aTF/DNA ratio improves sensitivity until aTF concentration approaches the aTF:DNA dissociation constant (14 pM), at which point leak becomes significant. **b.** The double equilibrium model predicts that further sensitivity improvements can be achieved if aTF:DNA and aTF:ligand dissociation constants are lowered from their literature values of 0.79 pM and 14 pM to 0.0079 pM and 0.14 pM, respectively. Compared with Figure 6a, significantly reduced levels of leak are observed at lower aTF concentrations. **c.** Dose response with tetracycline measured at 1 hr for both PSR and TMSD-only schemes. A 4.6-fold decrease in EC50 was observed with PSR. **d.** Application of PSR to zinc sensing with SmtB. Dose response with ZnSO4 measured after 1hr for both PSR and TMSD-only schemes. A 3.6-fold decrease in EC50 was observed with PSR. Data shown in c and d are n = 3 experimentally independent replicates, each plotted as a point with raw fluorescence standardized to MEF (µM FITC). Dotted curves represent Hill Equation fits (see Methods), with fit parameters in Source Data.

### PSR signal amplification can be used to increase biosensor sensitivity

Given that the model predicted enhanced sensitivity characterized by EC50, we next tested PSR with the tetracycline-sensing circuit. Using the previously optimized aTF and DNA template concentrations (Supplementary Figures 6, 7), we observed that the EC50 of the dose response with tetracycline was decreased by 4.6-fold with PSR (0.07 µM) compared to a TMSD- only system (0.32 µM), which is a 2.4-fold greater decrease than the 1.9-fold decrease predicted by the double equilibrium model (Figure 6c). Compared to the previously characterized EC50 for tetracycline sensing with TMSD circuits (1.03 µM) [28], we demonstrated a 14.7-fold reduction in EC50 due to the fact that the previously published work used a higher amount of DNA template and TetR than the TMSD-only controls in this study.

To test the generality of the system, we next applied it to zinc-sensing with the SmtB transcription factor. SmtB is the canonical zinc-sensing aTF from the ArsR/SmtB family of regulators that responds to metal ions [40]. Similar to TetR, SmtB is also an aTF that binds to its cognate operator sequence as a dimer, *smtO*, and can be de-repressed by zinc as well as Co(II) and Cd(II) [41]. Using a similar approach as for TetR, we first determined the DNA template concentration to use for PSR and TMSD-only systems and chose 1 nM of PSR DNA template and 25 nM of TMSD DNA template to use with circuits configured with *smtO* (Supplementary Figure 8). We next optimized [SmtB dimer]:[DNA] ratios to find the minimum amount of SmtB dimer needed to repress each system which was 0.25 µM (250X) for PSR and 0.625 µM (25X) for TMSD (Supplementary Figure 9). With the optimized component concentrations, we then performed a titration with ZnSO4 and found that the EC50 of the zinc-sensing biosensor could also be reduced with PSR (0.46 µM) by 3.6-fold compared to TMSD (1.66 µM) (Figure 6d), representing a 15.1-fold EC50 reduction in EC50 compared to previously published work (6.95 µM) which used a higher amount of DNA template and SmtB than the TMSD-only controls in this study [28].

Overall, these results showed that PSR signal amplification can be generalized and applied to biosensors based on both the TetR and SmtB/ArsR families of transcription factors, and that sensitivity enhancements can be achieved through PSR as hypothesized.

## Discussion

In this study, we present the development of polymerase strand recycling - a new concept in nucleic acid circuitry and DNA nanotechnology that uses T7 RNAP off-target transcription to create autocatalytic cycles of signal amplification of TMSD circuits. By using transcription to regenerate TMSD circuit inputs, the PSR process acts conceptually similarly to ‘fuel’ strands that regenerate inputs in other TMSD autocatalytic schemes [32]. However, by designing PSR to function in the context of *in vitro* transcription reactions, we show that PSR can be used as a signal amplification module for cell-free biosensing circuits that can sense RNA targets and small molecules, thus expanding the function of autocatalytic TMSD signal amplification. In this way, PSR is another type of information processing circuit system that can improve overall biosensor performance without having to perform protein engineering of the components [28].

The concept of PSR is inspired by previous work in DNA nanotechnology that developed nucleic acid signal amplification circuits such as catalytic hairpin assembly (CHA) [42, 43] and hybridization chain reaction (HCR) [44] which use TMSD to recycle a nucleic acid strand to enable an autocatalytic cycle of amplification. However, these existing circuits are not compatible with transcriptional biosensors due to exposed 3’ toeholds that cause T7 RNA Polymerase (RNAP) off-target transcription [28, 29]. PSR thus represents an advance of the fuel concept in two ways: (1) it allows fuel-driven signal amplification to be compatible with transcriptional circuitry, and (2) instead of using a nucleic acid fuel, it uses the RNA polymerase enzyme. The latter concept is important, as in PSR the fuel-driven reactions are not limited by the amount of specific fuel strand supplied in the reactions, but rather could continue as long as T7 transcription was supported by the reaction. We therefore anticipate creative uses of the PSR concept in a range of DNA nanotechnology circuits and applications.

An intriguing feature of signal amplification is that it can potentially be used to enhance biosensor sensitivity. Specifically, a double equilibrium model of aTF:DNA and aTF:ligand binding interactions predicts that decreasing aTF and DNA template concentration can improve biosensor sensitivity characterized by EC50 (Figure 6a). Signal amplification enables this prediction to be realized in practice because it lowers the concentrations of aTF and DNA template required to enact sensing function without increasing sensor leak. Theoretically, we can lower the aTF concentration required to sufficiently repress transcription down to near its KD, where KD is the aTF:DNA dissociation constant. For the TetR system, this dissociation constant is approximately 0.014 nM, though this limit may be unattainable in practice if stochastic effects begin to dominate at low concentrations. In our present study, we show that in the case of tetracycline sensing with the TetR aTF, PSR can provide a 4.6-fold reduction in EC50 and in the case of zinc sensing with the SmtB aTF, PSR can provide a 3.6-fold reduction in EC50 compared to TMSD biosensing circuits without signal amplification (Figure 6c-d). These results overall support the model, but do not exactly match model predictions. This could be caused by several factors, including the potential for PSR to generate and amplify fluorescent signal from low amounts of unbound DNA template which causes a non-linear relationship between the amount of unbound DNA template and observed fluorescence output, potential problems with aTF quality causing a fraction of the aTF population to be non-functional, and discrepancies between the dissociation constants used in the model and those apparent in the actual system. With further improvements in aTF purification strategies that can generate high quality proteins, and further engineering of the system, it may be possible to get closer to the theoretical limit, which could greatly improve the performance of cell-free biosensors and allow them to meet various regulatory and recommended detection thresholds for a range of important contaminants [45].

T7 RNAP off-target transcription from 3’ toeholds is typically presented as a design limitation in DNA nanotechnology circuits [28, 29]. Here we were able to turn this ‘bug’ into a ‘feature’ by developing design rules to control and leverage this behavior to act as a fuel that recycles input nucleic acid strands. One challenge we encountered was the inhibiting effects of DNA-RNA duplexes on T7 RNAP off-target transcription (Figure 2). Based on previous reports showing that DNA-RNA duplexes appear to adopt an overall A-form while DNA-DNA duplexes adopt B-form helices, we speculate that the DNA-RNA duplex conflicts with the polymerase structure and inhibits 3’ toehold off-target transcription [46]. We were ultimately able to address this challenge by implementing an intermediate fuel gate design that in effect takes an input RNA and converts it to a DNA strand that can then participate in strand recycling (Figure 3).

For the most part, the design of PSR circuits follows simple rules of designing a fuel gate with an overlap of 18- to 22-bp between RecycleD and a 2’-O-methyl modified strand to sequester the RNA input, and designing a signal gate that can be strand invaded by RecycleD to form a new duplex with a 3’-toehold for T7 RNAP off-target transcription (Supplementary Figures 3, 5). Using these rules, we were able to design a modular system that works with transcription, and that can sense different compounds with aTFs by changing the aTF operator site in the transcription template (Figure 6). Applying these concepts to direct detection of RNAs was also possible by designing the sequence of the fuel gate to be strand invaded by an RNA input sequence while maintaining a ∼20-bp overlap between RecycleD and the 2’-O-methyl modified strand. This worked well in the context of detecting miRNA sequences, where we were able to design systems that can detect *in vitro* synthesized versions of miR-3185, miR-134-3p and miR-642a-5p with high degrees of specificity and LoDs on the order of 5 – 250 nM.

One limitation of the PSR system is the cost of the chemically modified gate components. We found that it was important to introduce 2’-O-methyl modified strand to form the duplex with RecycleD in the fuel gate to reduce leak in the absence of input and improve the biosensor dynamic range. While each set of gate designs with these modifications currently cost ∼100 USD, we found we could still perform hundreds of reactions with each batch of oligos. Fortunately, the PSR system is highly modular, allowing us to use the same set of fuel gate and signal gate for different sensing targets to justify the cost of oligos.

We believe the PSR circuit can be interfaced with other molecular computation circuits to expand capabilities of cell-free systems. For example, PSR could serve as an amplified output for logic gates [3, 28, 47], genelets [48, 49], co-transcriptionally activated RNA gates [50, 51], or other signal processing circuits to control complex computations in cell-free systems [52]. We also envision using PSR to amplify RNA transcriptional signals from other polymerases that are less efficient than T7 RNAP, such as *E. coli* RNAP, to enable robust signal generation.

Overall, we believe PSR will serve as an important component in an expanding toolbox of cell-free nucleic acid circuitry that can be used to enhance biosensing reaction performance. In this way they enhance the many strong features of cell-free biosensors including their accessibility, relatively low cost, modularity and ability to operate in field conditions [2, 5, 16, 28]. In addition, we anticipate other creative applications of the PSR concept in a range of biotechnologies, including TMSD circuits, where signal amplification and genetic circuits are important for function.

## Materials and methods

### Strains and growth medium

*Escherichia coli* strain K12 (NEB Turbo Competent *E. coli*, New England Biolabs, catalog no. C2984) was used for routine cloning. *E. coli* strain Rosetta 2(DE3)pLysS (Novagen, catalog no. 71401) was used for recombinant protein expression. Luria broth supplemented with the appropriate antibiotic(s) (100 µg ml^−1^ carbenicillin, 100 µg ml^−1^ kanamycin and/or 34 µg ml^−1^ chloramphenicol) was used as the growth media.

### DNA gate preparation

DNA fuel gates and signal gates used in this study were synthesized by Integrated DNA technologies as HPLC purified and chemically modified oligos (Supplementary Data). Gates were generated by denaturing complementary strands at 95° C for 5 min and slow cooling (– 0.1 °C s^−1^) to room temperature in annealing buffer (50 mM Tris-HCl, pH 8.0, 10 mM MgCl2). DNA fuel gates and signal gates were stored at 4°C until use.

### Plasmids and genetic parts assembly

DNA oligonucleotides for cloning and sequencing were synthesized by Integrated DNA Technologies. Genes encoding aTFs were synthesized either as gBlocks (Integrated DNA Technologies) or gene fragments (Twist Bioscience). Protein expression plasmids were cloned using Gibson Assembly (NEB Gibson Assembly Master Mix, New England Biolabs, catalog no. E2611) into a pET-28c plasmid backbone and were designed to overexpress recombinant proteins as C-terminus His-tagged fusions. A construct for expressing SmtB additionally incorporated a recognition sequence for cleavage and removal of the His-tag using TEV protease. Gibson assembled constructs were transformed into NEB Turbo cells, and isolated colonies were purified for plasmid DNA (QIAprep Spin Miniprep Kit, Qiagen, catalog no. 27106). Plasmid sequences were verified with Sanger DNA sequencing (Quintara Biosciences) using the primers listed in Supplementary Data.

All transcription templates were generated using PCR amplification (Phusion High-Fidelity PCR Kit, New England Biolabs, catalog no. E0553) of an oligo that includes a T7 RNAP promoter, an optional aTF operator site, and the InvadeR or RecycleR coding sequence. All oligos and primer sets used in this study are listed in Supplementary Data. Here, we define the T7 RNAP promoter as a minimal 17-bp sequence (TAATACGACTCACTATA) excluding the first G that is transcribed. The PCR-amplified templates were purified (QIAquick PCR purification kit, Qiagen, catalog no. 28106) and verified for the presence of a single DNA band of expected size on a 2% Tris-Acetate-EDTA–agarose gel. Concentrations of all DNA templates were determined using the Qubit dsDNA BR Assay Kit (Invitrogen, catalog no. Q32853).

All plasmids and DNA templates were stored at 4 °C until use. A spreadsheet listing the sequences and the Addgene accession numbers of all plasmids and oligos used or generated in this study are listed in Supplementary Data.

### RNA expression and purification

RecycleR used for Figure 2 was first expressed by an overnight IVT at 37 °C from a transcription template encoding a *cis*-cleaving hepatitis D ribozyme on the 3′-end of the RecycleR sequence with the following components: IVT buffer (40 mM Tris–HCl pH 8, 8 mM MgCl2, 10 mM dithiothreitol, 20 mM NaCl and 2 mM spermidine), 11.4 mM NTPs pH 7.5, 0.3 units (U) of thermostable inorganic pyrophosphatase (New England Biolabs, catalog no. M0296S), 100 nM transcription template, 50 ng of T7 RNAP and MilliQ ultrapure H2O to a total volume of 500 µl. The overnight IVT reactions were then ethanol-precipitated and purified by resolving them on a 20% urea–PAGE–TBE gel, isolating the band of expected size (26–29 nucleotides) and eluting at 4 °C overnight in MilliQ ultrapure H2O. The eluted InvadeR and RecycleR variants were ethanol-precipitated, resuspended in MilliQ ultrapure H2O, quantified using the Qubit RNA BR Assay Kit (Invitrogen, catalog no. Q10211) and stored at –20 °C until use. The hepatitis D ribozyme sequence used can be found in Supplementary Data. miRNA used for miRNA sensing experiments were synthesized and PAGE purified by Integrated DNA Technologies.

### aTF expression and purification

aTFs were expressed and purified as previously described [2]. Briefly, sequence-verified pET- 28c plasmids were transformed into the Rosetta 2(DE3)pLysS *E. coli* strain. Cell cultures (1–2 L) were grown in Luria broth at 37 °C, induced with 0.5 mM of isopropyl-β-D-thiogalactoside at an optical density (600 nm) of ∼0.5 and grown for a further 4 h at 37 °C. Cultures were then pelleted by centrifugation and were either stored at –80 °C or resuspended in lysis buffer (10 mM Tris–HCl pH 8, 500 mM NaCl, 1 mM Tris(2-carboxyethyl)phosphine (TCEP) and protease inhibitor (Complete EDTA-free Protease Inhibitor Cocktail, Roche)) for purification. Resuspended cells were then lysed on ice through ultrasonication, and insoluble materials were removed by centrifugation. Clarified supernatant containing TetR was then purified using His-tag affinity chromatography with a Ni-NTA column (HisTrap FF 5 ml column, GE Healthcare Life Sciences) followed by size-exclusion chromatography (Superdex HiLoad 26/600 200 pg column, GE Healthcare Life Sciences) using an AKTAxpress fast protein liquid chromatography system. For SmtB expression and purification cultures and induction were performed in the same fashion as TetR, pellets were resuspended with a previously degassed lysis buffer (25mM MES, 750mM NaCl 1mM EDTA, pH 6, 2mM TCEP, 0.1mM PMSF) and then lysed via ultrasonication. Lysates were then centrifuged, and the supernatant treated with 0.015 %V/V of Poly(ethyleneimine) solution. The suspension was again centrifuged, and the supernatant taken to 70% ammonium sulfate. The suspension was centrifuged, and the pellet resuspended in lysis buffer with 75mM NaCl, for subsequent dialysis with the same buffer in a 10kDa dialysis bag.

The dialysis product was centrifuged, and the supernatant loaded into a SP cation exchange column to be eluted with a gradient of high salt lysis buffer. The fractions containing protein were pooled and treated for 16 hours with TEV protease, then loaded into a Superdex 75 size exclusion column equilibrated with S75 buffer (25mM Tris, 200mM NaCl, pH 8, degassed, 2mM TCEP). Fractions corresponding with the protein’s weight were pooled and concentrated.

Protein concentrations were measured via the Qubit Protein Assay Kit, and protein purity and size were verified via SDS-PAGE gel. Purified proteins were stored at -20 °C. The eluted fractions from the fast protein liquid chromatography for TetR were concentrated and buffer exchanged (25 mM Tris–HCl, 100 mM NaCl, 1 mM TCEP, 50% glycerol v/v) using centrifugal filtration (Amicon Ultra-0.5, Millipore Sigma). Protein concentrations were determined using the Qubit Protein Assay Kit (Invitrogen, catalog no. Q33212). The purity and size of the proteins were validated on an SDS–PAGE gel (Mini-PROTEAN TGX and Mini-TETRA cell, Bio-Rad). Purified proteins were stored at –20 °C.

### PSR reactions

PSR reactions and miRNA-sensing PSR reactions were set up by adding the following components listed at their final concentration: IVT buffer (40 mM Tris–HCl pH 8, 8 mM MgCl2, 10 mM dithiothreitol, 20 mM NaCl and 2 mM spermidine), 11.4 mM NTPs pH 7.5, 0.3 U of thermostable inorganic pyrophosphatase (New England Biolabs, catalog no. M0296S), transcription template, DNA gate(s) and MilliQ ultrapure H2O to a total volume of 20 µL.

Regulated PSR reactions additionally included a purified aTF at the indicated concentration and were incubated at 37 °C for ∼10 min.

Immediately before plate reader measurements, 2 ng of T7 RNAP and, optionally, a ligand or purified miRNA at the indicated concentration were added to the reaction. Reactions were then characterized on a plate reader as described in ‘Plate reader quantification and micromolar equivalent fluorescein standardization’.

### Plate reader quantification and micromolar equivalent fluorescein standardization

A National Institute of Standards and Technology traceable standard (Invitrogen, catalog no. F36915) was used to convert arbitrary fluorescence measurements to micromolar equivalent fluorescein (MEF). Serial dilutions from a 50 µM stock were prepared in 100 mM sodium borate buffer at pH 9.5, including a 100 mM sodium borate buffer blank (total of 12 samples). For each concentration, three replicates of samples were created in batches of three, and fluorescence values were read at an excitation wavelength of 490 nm and emission wavelength of 525 nm for 6-FAM (fluorescein)-activated fluorescence (Synergy H1, BioTek Gen5 v.2.04). Fluorescence values for a fluorescein concentration in which a single replicate saturated the plate reader were excluded from the analysis. The remaining replicates (three per sample) were then averaged at each fluorescein concentration, and the average fluorescence value of the blank was subtracted from all values. Linear regression was then performed for concentrations within the linear range of fluorescence (0–3.125 µM fluorescein) between the measured fluorescence values in arbitrary units and the concentration of fluorescein to identify the conversion factor. For each plate reader, excitation, emission and gain setting, we found a linear conversion factor (setting the y intercept to 0) that was used to correlate arbitrary fluorescence values to MEF (Supplementary Figure 1).

For reaction characterization, 19 µL of reactions were loaded onto a 384-well optically clear, flat-bottom plate using a multichannel pipette, covered with a plate seal and measured on a plate reader (Synergy H1, BioTek Gen5 v.2.04). Kinetic analysis of 6-FAM (fluorescein)-activated fluorescence was performed by reading the plate at 1-min intervals with excitation and emission wavelengths of 490 and 525 nm, respectively, for 1 hr (ligand-sensing reactions) or 2 hrs (miRNA-sensing reactions) at 37 °C. Arbitrary fluorescence values were then converted to MEF by dividing by the appropriate calibration conversion factor.

### Hill equation fits

Where indicated, data were fit to the Hill equation with the following functional form

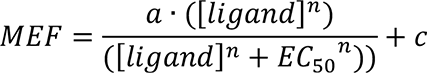

Where [ligand] denotes ligand concentration, *c* represents the response with no ligand, *a* represents the maximum response, and *n* describes the cooperativity. Fitting was performed on individual replicate datasets using DataGraph 5.2 by starting with a = 1.5, EC50 = 1, c = 1, n = 1, then optimized for exact parameters with DataGraph 5.2. Curves were generated from fitting the average of the replicates for plotting. For TetR and SmtB dose response curves, EC50 values were obtained from the fits of the average of the replicates.

## STATISTICS AND REPRODUCIBILITY

The number of replicates and types of replicates performed are described in the legend of each figure. Individual data points are shown, and where relevant, the average ± s.d. is shown; this information is provided in each figure legend. The type of statistical analysis performed in Figures 2b, 3b, 4b, and 4c is described in the legend to each figure. Exact p-values along with degrees from the statistical analysis can be found in Source Data.

## DATA AVAILABILITY

All data presented in this paper are available as Source Data and as Supplementary Data. Both plasmids used in this paper are available in Addgene with the identifiers 140371 and 140395.

Source Data are provided with this paper.

## CODE AVAILABILITY

The Jupyter Notebook files with the Python codes used in Figure 6a and Figure 6b are provided as Supplementary Data, and the double equilibrium model used in this paper is described in the Supplementary Information.

## AUTHOR CONTRIBUTIONS

Y.L., J.K.J. and J.B.L. designed the study. Y.L. and J.B.L. analyzed the data. Y.L., M.V.D. and T.L. conducted the research. Y.L, J.K.J., M.V.D., D.A.C. and J.B.L. developed the methodology. Y.L. and J.B.L. undertook visualization of the data. T.L. developed the software. Y.L and J.B.L. curated the data. J.B.L. acquired funding for the study. Y.L and T.L. validated the results. Y.L. and J.B.L. managed and coordinated the study. Y.L., D.A.C. and J.B.L. supervised the research. Y.L., T.L. and J.B.L. wrote the manuscript. All authors edited the manuscript.

## Supporting information

All Source Data

Jupyter Notebook Code for Supplementary Note

## ACKNOWLEDEGMENTS

We thank C. Knopp (Northwestern University) and A. Moreno (Northwestern University) for managing the experimental reagents and equipment used in this study; S. Schaffter (National Institute of Standards and Technology) for helpful discussion on TMSD circuits and T7 RNAP activity; S. O. Kelley (Northwestern University), E.H. Sargent (Northwestern University), A. D. Ellington (University of Texas at Austin), M. C. Jewett (Stanford University), and J. L. Chavez (AFRL) for helpful discussions on miRNA target selection. Y.L. was supported by the National Institutes of Health Training Grant (T32GM008449) through Northwestern University’s Biotechnology Training Program and the Ryan Fellowship and the International Institute for Nanotechnology at Northwestern University. T.L was supported by the National Science Foundation Synthesizing Biology Across Scales NRT training program (2021900). J.K.J. was supported by Northwestern University’s Graduate School Cluster in Biotechnology, System, and Synthetic Biology, which is affiliated with the Biotechnology Training Program, the Ryan Fellowship and the McCormick School of Engineering Terminal Year Fellowship. This work was also supported by Army Contracting Command (W52P1J-21-9-3023) and Defense Advanced Research Projects Agency (DARPA) (N660012324041), and the National Science Foundation (2310382) to J.B.L. The views, opinions, and/or findings expressed are those of the authors and should not be interpreted as representing the official views or policies of the Department of Defense, the National Science Foundation or the U.S. Government. D.A.C. and M.V.D. were supported by Bunge & Born, Argentina; Williams Foundations; MinCyT Argentina (PICT 2022- ET-09-00049 and B81 CYTCH), D.A.C is a staff member from CONICET, Argentina; M.V.D. is supported by a fellowship provided by CONICET, Argentina.

