## Supplementary Information for "Engineering a cell-free biosensor signal amplification circuit with polymerase strand recycling"

Yueyi Li[1,2], Tyler Lucci[1,2], Matias Villarruel Dujovne[3], Jaeyoung Kirsten Jung[1,2], Daiana A. Capdevila[3], Julius B. Lucks[1,2,4,5,6†]

[1]Department of Chemical and Biological Engineering, Northwestern University, Evanston, Illinois 60208, USA
[2]Center for Synthetic Biology, Northwestern University, Evanston, Illinois 60208, USA
[3]Fundación Instituto Leloir, Buenos Aires, Argentina
[4]Interdiscipinary Biological Sciences Graduate Program, Northwestern University, Evanston, Illinois 60208, USA
[5]Center for Water Research, Northwestern University, Evanston, Illinois 60208, USA
[6]Center for Engineering Sustainability and Resilience, Northwestern University, Evanston, Illinois 60208, USA

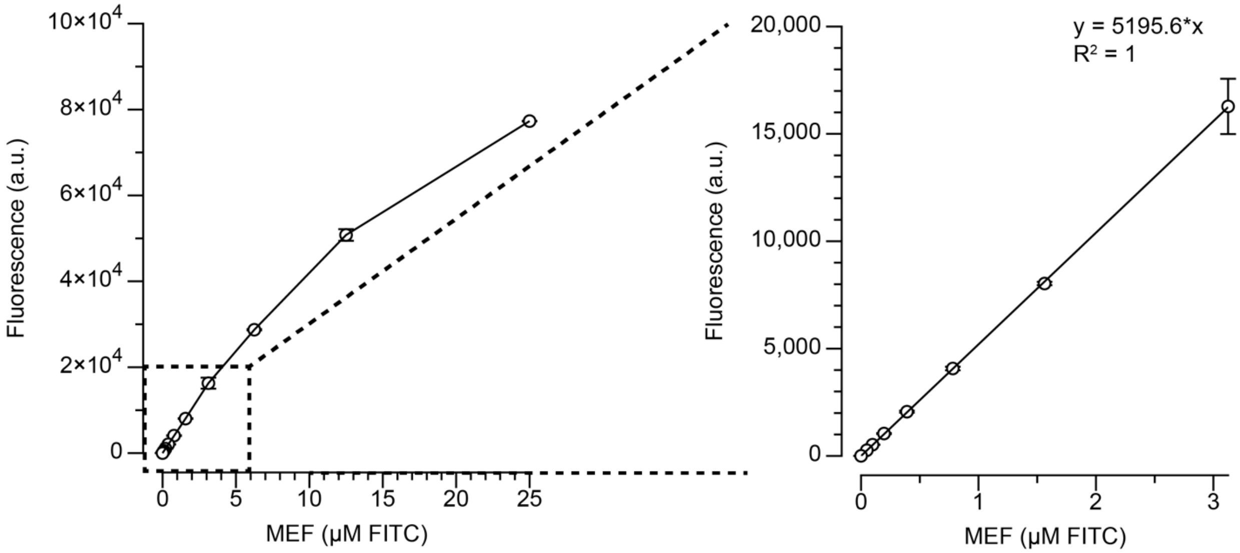


**SI Figure 1. Micromolar Equivalent Fluorescein (MEF) standardization.** Arbitrary units of fluorescence were standardized to µM equivalent fluorescein (µM FITC) using a NIST traceable standard (see Methods). In the representative example shown here, a dilution series of FITC standard was prepared in buffer (100 mM sodium borate, pH 9.5) and measured on a plate reader using the same settings for measuring 6-FAM signal (490 nm excitation, 525 nm emission). The resulting curve, calculated over the linear range of 0–3.125 µM, was then used to standardize fluorescence measured from PSR and TMSD reactions. The standard curve was generated at regular intervals for each plate reader and each measurement setting. Data shown are for n=3 experimentally independent replicates for each concentration. Error bars indicate standard deviation computed over n=3 replicates.

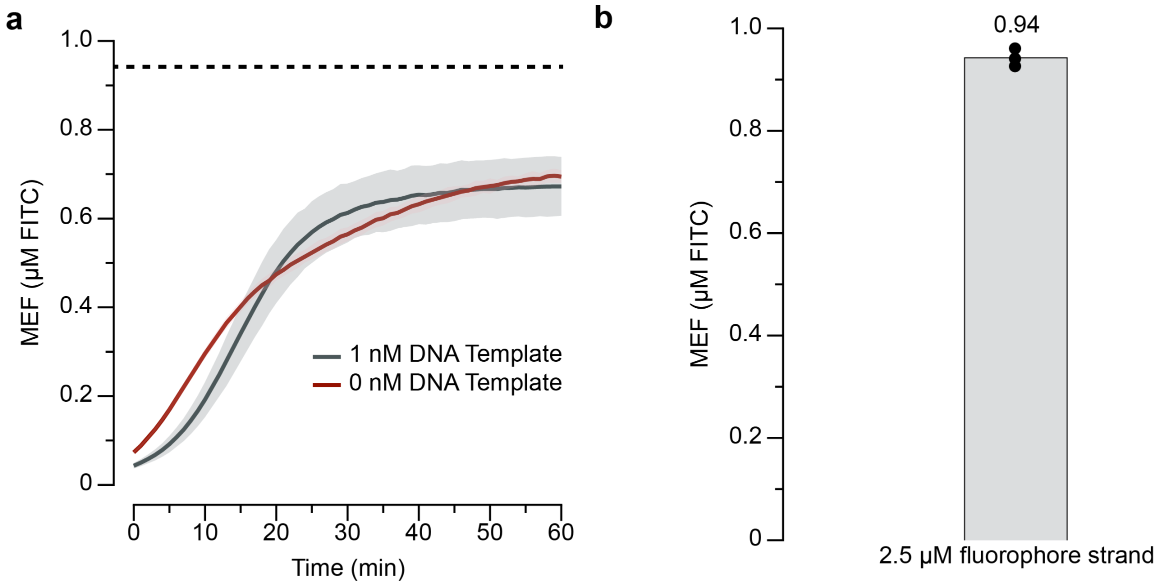


**SI Figure 2. T7 RNAP can activate fluorescence signal from fuel gate and signal gate even in the absence of DNA template. a.** Both 1 nM DNA template and 0 nM DNA template activated fluorescent signal after 1 hr with 1.5 µM fuel gate and 2.5 µM signal gate. A dashed line indicates the fluorescence level of 2.5 µM of pure fluoprophore strand in IVT buffer. Solid lines are averages over n = 3 experimentally independent replicates. Shading indicates the average of the replicates ± s.d. **b.** Measured MEF (µM FITC) of 2.5 µM of fluorescent fluorophore strand in IVT buffer. Data shown are n = 3 experimentally independent replicates each plotted as a point with raw fluorescence standardized to MEF (µM FITC). Bar heights represent the average over these replicates.

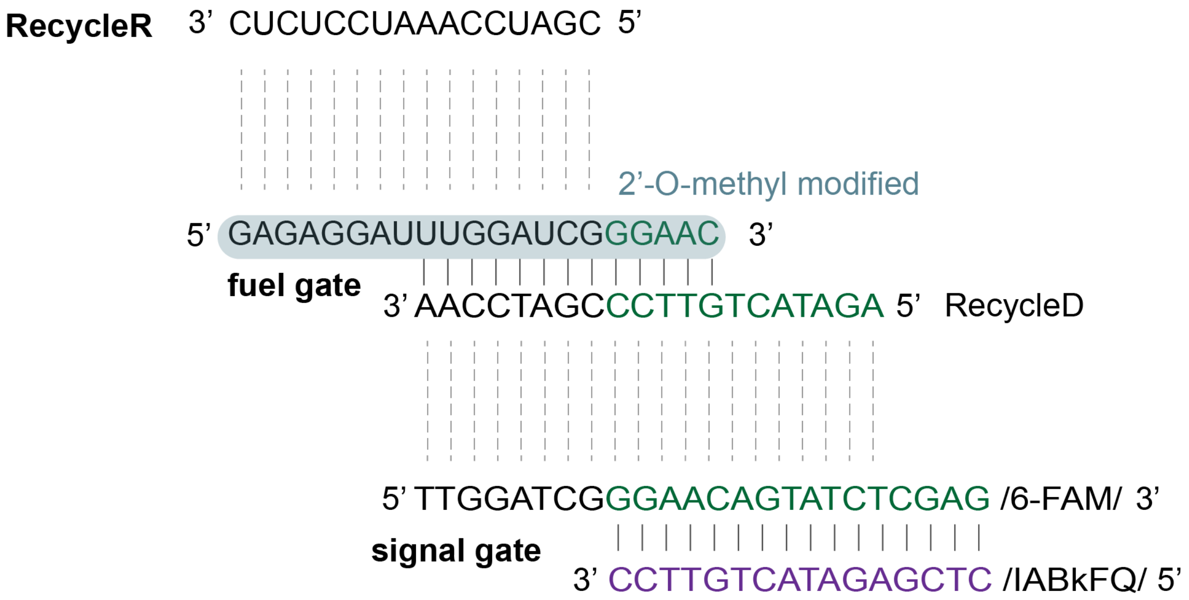


**SI Figure 3. Design elements in a PSR circuit.** Sequences for RecycleR, fuel gate, and signal gate. The fuel gate can be strand invaded by RecycleR. It consists of a 2'-O-methyl modified strand and a DNA strand, RecycleD. The DNA signal gate has a fluorophore strand and a quencher strand.

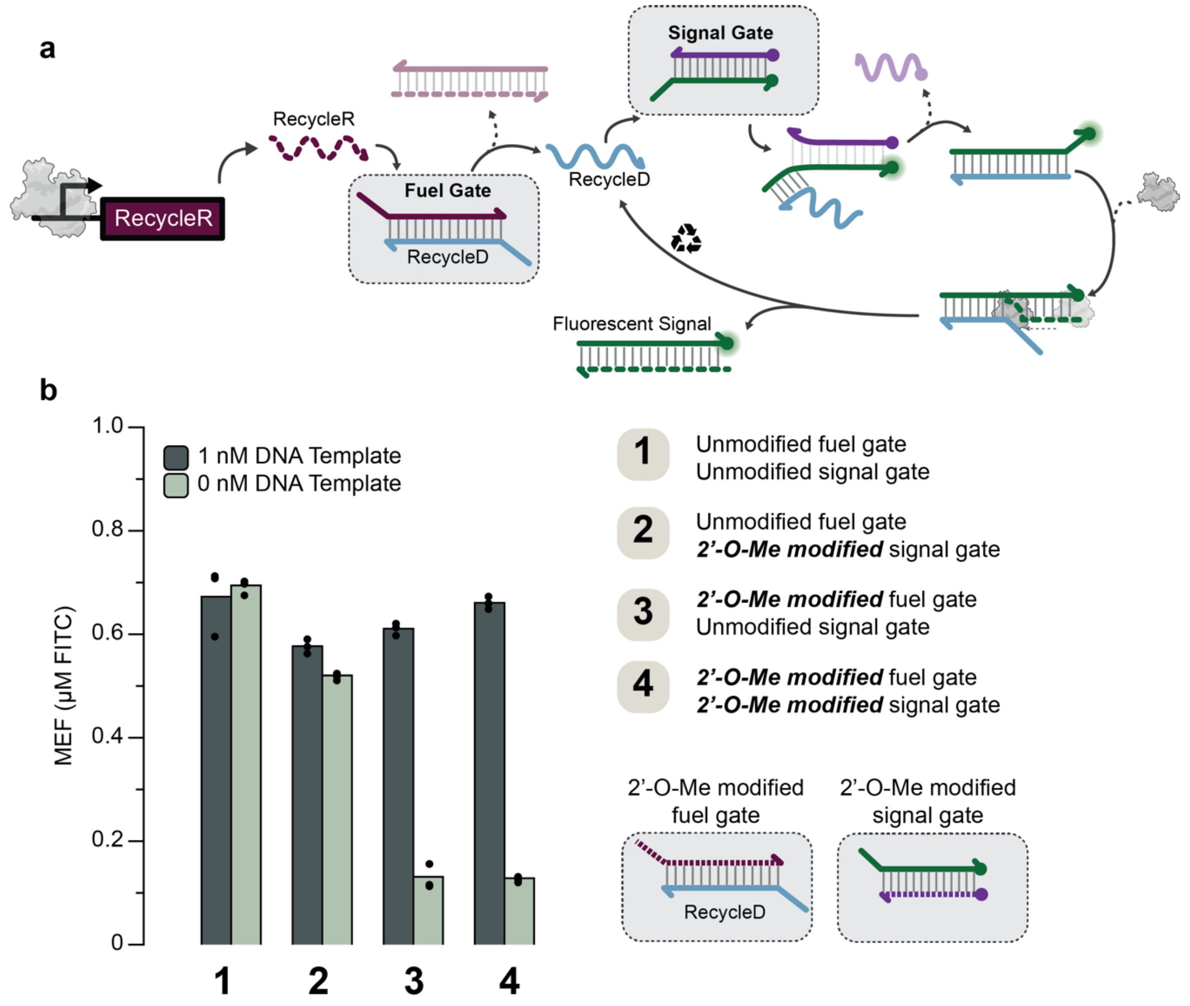


**SI Figure 4. 2'-O-Me modification significantly reduces the amount of leak in PSR circuits.** **a.** Circuit scheme after including the fuel gate concept. **b.** Fluorescence was characterized for different combinations of unmodified and 2'-O-Me modified fuel and signal gate in the presence of 1.5 µM fuel gate, 2.5 µM signal gate, 2 ng T7 RNAP, and the indicated amount of DNA template encoding RecycleR. Dotted lines represent 2'-O-Me modification. Data shown as points are for n = 3 experimentally independent replicates. Bar heights represent the average over these replicates.

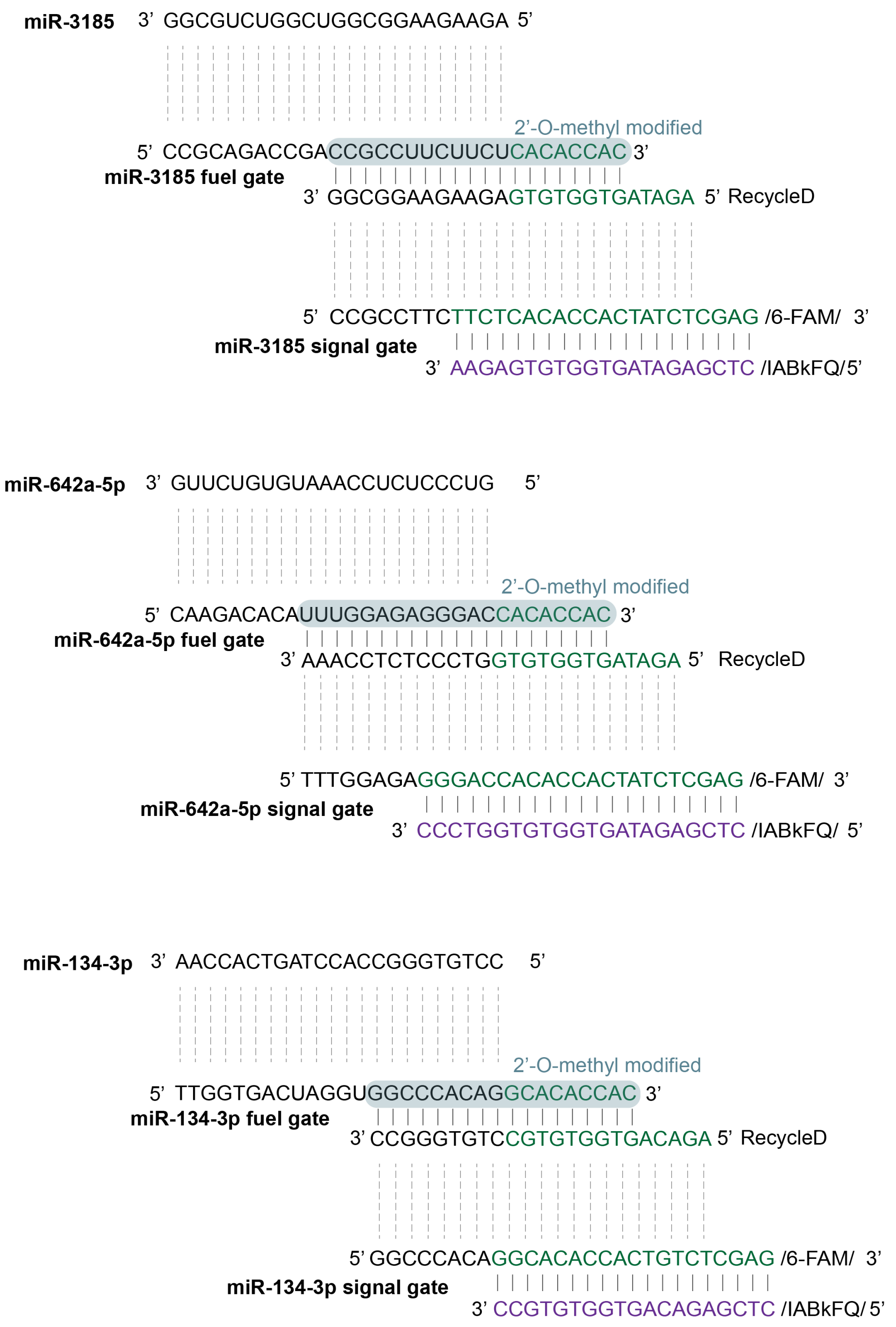


**SI Figure 5. Design elements in PSR circuits to detect RNA inputs.** Sequences for the miRNAs, and their respective fuel gates and signal gates. The fuel gates can be strand invaded by miRNAs. They consist of a 2'-O-methyl modified strand and an unmodified RecycleD DNA strand. The DNA signal gates have a fluorophore strand and a quencher strand.

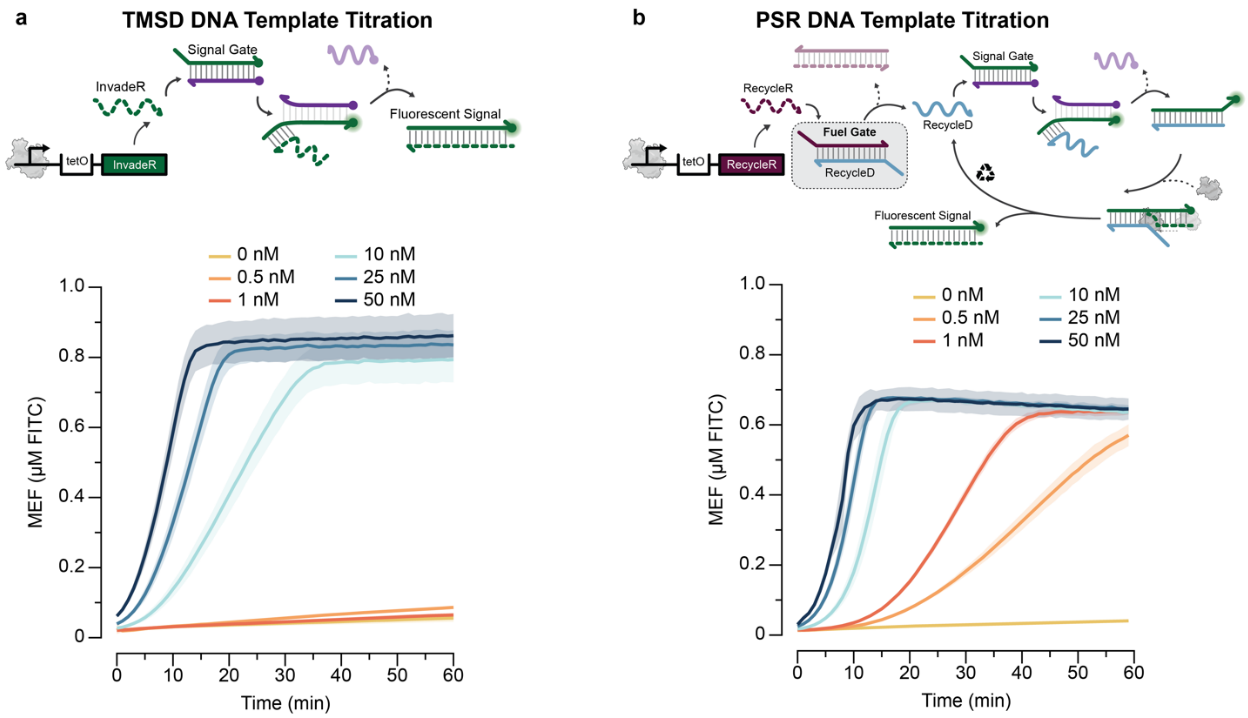


**SI Figure 6. Titrating DNA templates containing the *tetO* sequence to compare TMSD signal generation to signal generation with PSR.** Time course trajectories of fluorescence from **a.** TMSD circuits and **b.** PSR circuits with various amounts of input DNA template added. TMSD circuits contained 2 ng of T7 RNAP, 2.5 µM DNA signal gate and various amounts of input DNA template. PSR circuits contained 2 ng of T7 RNAP, 1.5 µM fuel gate, 2.5 µM DNA signal gate and various amounts of input DNA template. Data shown are from n = 3 experimentally independent replicates each plotted as a line. Shading indicates the average of the replicates ± s.d. Data in b repeated from Figure 5b for comparison.

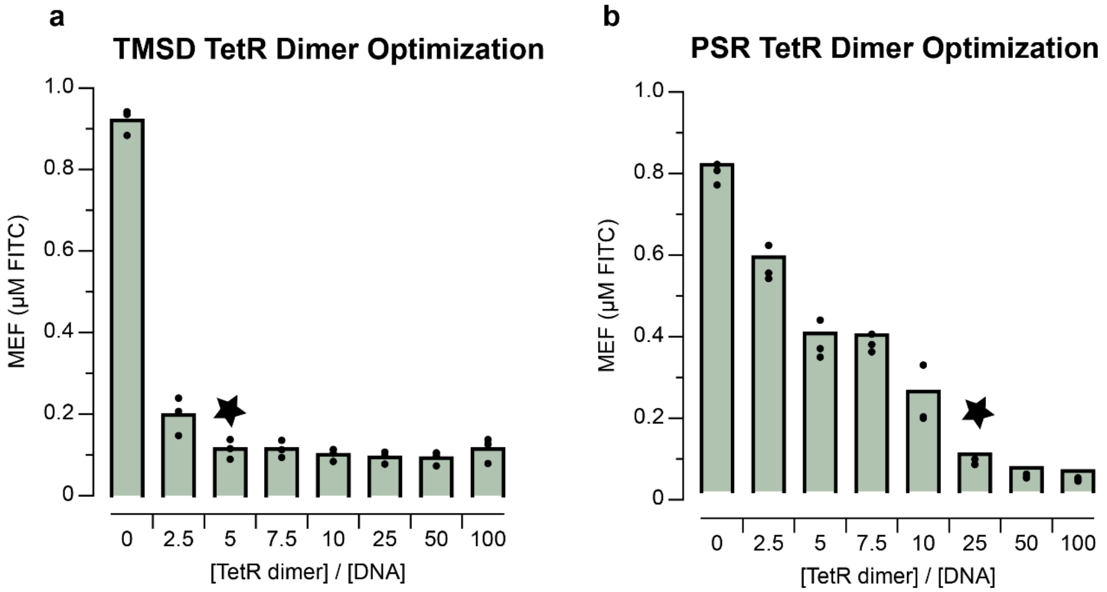


**SI Figure 7. Titrating [TetR dimer] at a fixed [DNA] to identify the amount of TetR needed to fully repress transcription for TMSD and PSR biosensing circuits. a.** 5X [TetR dimer] to [DNA] ratio was shown to fully repress transcription in TMSD biosensing circuits (star), corresponding to 50 nM TetR dimer to 10 nM TMSD DNA template. **b**. 25X [TetR dimer] to [DNA] ratio was shown to fully repress transcription in PSR biosensing circuits (star), which was 12.5 nM TetR dimer to 0.5 nM of PSR DNA template. Data in a and b shown are for n = 3 experimentally independent replicates shown as points. Bar heights represent the average over these replicates. Data in b repeated from Figure 5c for comparison.

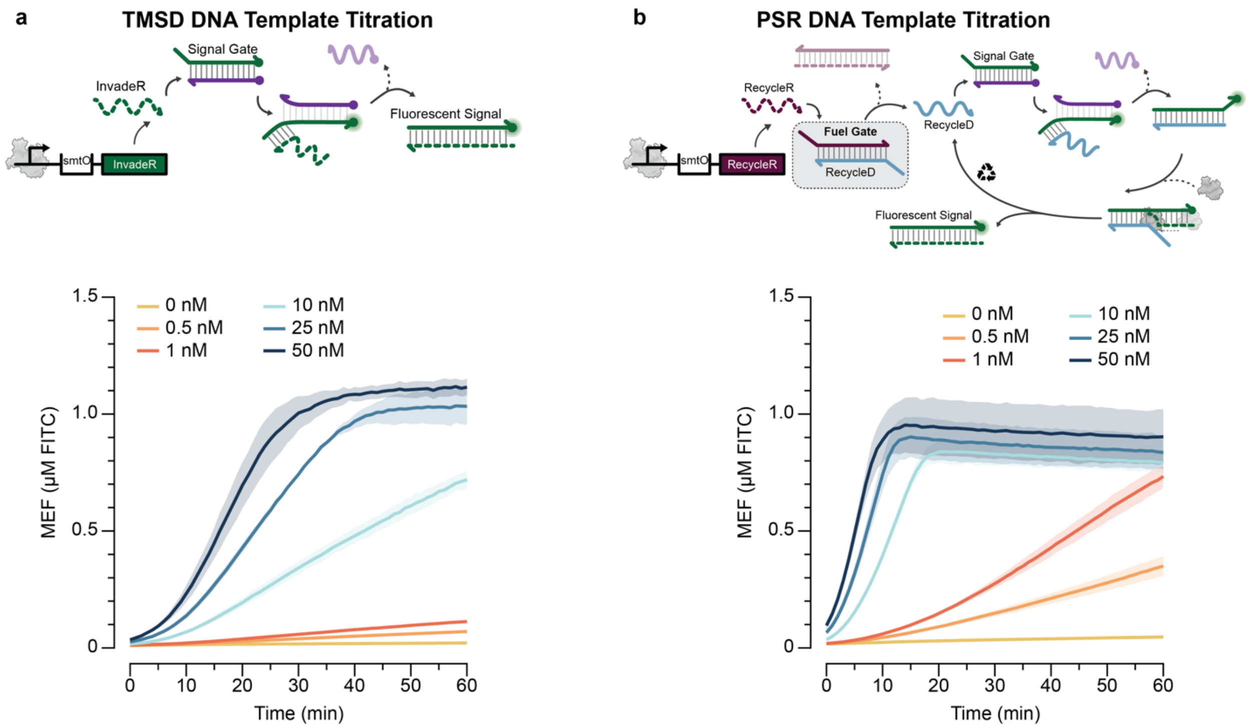


**SI Figure 8. Titrating DNA templates containing the *smtO* sequence to compare TMSD signal generation to signal generation with PSR.** Time course trajectories of fluorescence from **a.** TMSD circuits and **b.** PSR circuits. TMSD circuits contained 2 ng of T7 RNAP, 2.5 µM DNA signal gate and various amounts of input DNA template. PSR circuits contained 2 ng of T7 RNAP, 1.5 µM fuel gate, 2.5 µM DNA signal gate and various amounts of input DNA template. Data shown are from n = 3 experimentally independent replicates, with lines representing the average. Shading indicates the average of the replicates ± s.d.

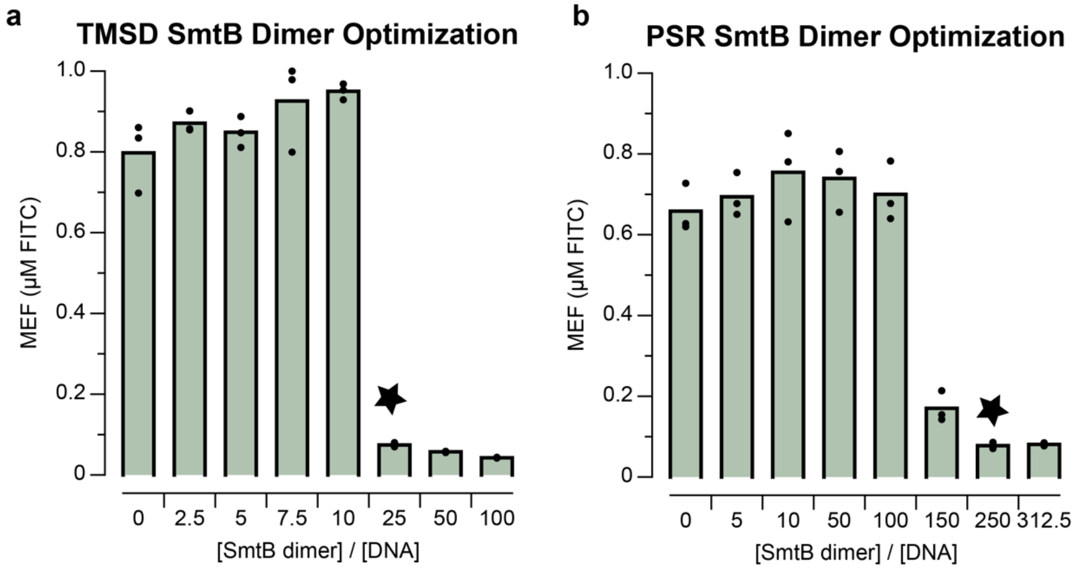


**SI Figure 9. Titrating [SmtB dimer] at a fixed [DNA] to identify the amount of SmtB to fully repress transcription for TMSD and PSR biosensing circuits. a.** 25X [SmtB dimer] to [DNA] ratio was shown to fully repress transcription in TMSD biosensing circuits (star), corresponding to 625 nM SmtB dimer to 25 nM TMSD DNA template. **b**. 250X [SmtB dimer] to [DNA] ratio was shown to fully repress transcription in PSR biosensing circuits (star), which was 250 nM SmtB dimer to 1 nM of PSR DNA template. Data in a and b shown are for n = 3 experimentally independent replicates shown as points. Bar heights represent the average over these replicates.

**Supplementary Note. A double equilibrium model predicts enhancement of biosensor sensitivity when less aTF is included.**

To predict expected transcription levels for different DNA/aTF/ligand concentrations, we developed a simple double equilibrium model describing the behavior of aTF:ligand and aTF:DNA binding interactions. The model takes as input DNA template concentration, aTF concentration, ligand concentration, DNA/aTF dissociation constant, and ligand/aTF dissociation constant, and solves for the fraction of DNA template without aTF bound, which relates to expected transcription level. The model contains five equations (species balances and dissociation equilibria) and five unknowns (species concentrations). Using substitution, the five equations can be reduced to a single nonlinear equation containing a single unknown, which can be solved numerically. The remaining unknowns can then be solved for via back substitution.

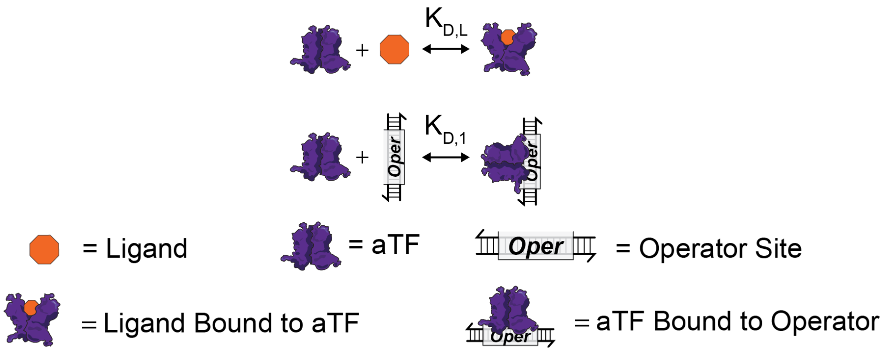


**SI Figure 10. Double equilibrium model of transcription factor binding to DNA and target compound.** The double equilibrium model considers association/dissociation of ligand-aTF (top) and of aTF-operator site (bottom). Species are indicated with schematics. aTF refers to the functional aTF unit which could be a multimer for some mechanisms.

**SI Table 1: Model notation**

| Abbreviation | Description |
|---|---|
| $\mathrm{K_{D,L}}$ | Dissociation constant for ligand and aTF |
| $\mathrm{K_{D,1}}$ | Dissociation constant for aTF and operator site on DNA |
| $[\mathrm{DNA}]_\mathrm{T}$ | Total DNA template concentration in solution |
| $[\mathrm{L}]_\mathrm{T}$ | Total ligand concentration in solution |
| $[\mathrm{aTF}]_\mathrm{T}$ | Total aTF concentration in solution |
| $[\mathrm{DNA}]$ | Concentration of DNA template in solution without aTF bound |
| $[\mathrm{aTF}]$ | Concentration of aTF in solution not bound to ligand or DNA |
| $[\mathrm{L} \cdot \mathrm{aTF}]$ | Concentration of aTF in solution bound to ligand |
| $[\mathrm{DNA} \cdot \mathrm{aTF}]$ | Concentration of aTF in solution bound to DNA |

<u>Equations:</u>

The definitions of the equilibrium dissociation constants are as follows:

1) $\mathrm{K_{D,L}} = \frac{[\mathrm{L}][\mathrm{aTF}]}{[\mathrm{L \cdot aTF}]}$

2) $K_{D,1} = \frac{[DNA][aTF]}{[DNA \cdot aTF]}$

The species conservation equations for DNA, ligand, and aTF are as follows:

3) $[DNA]_T = [DNA] + [DNA \cdot aTF]$
4) $[L]_T = [L] + [L \cdot aTF]$
5) $[aTF]_T = [aTF] + [L \cdot aTF] + [DNA \cdot aTF]$

<u>Solving the double equilibrium:</u>

We assume that the aTF can bind to either the DNA template or the ligand, but not both simultaneously. For a given experimental condition (specified by the input concentration of each molecular species), we assume that the fraction of DNA unbound by aTF is proportional to the transcription output signal generated. Therefore, the goal of the double equilibrium analysis is to solve for $[DNA]/[DNA]_T$ as a function of $[aTF]_T$ and $[L]_T$.

From (2) and (3):

$$[DNA] = K_{D,1}\frac{[DNA \cdot aTF]}{[aTF]} = K_{D,1}\frac{[DNA]_T - [DNA]}{[aTF]}$$

Rearranging gives:

$$[DNA]\left(1 + \frac{K_{D,1}}{[aTF]}\right) = K_{D,1}\frac{[DNA]_T}{[aTF]}$$

$[DNA]/[DNA]_T$ can then be written as a function of $[aTF]$:

6) $f = \frac{[DNA]}{[DNA]_T} = \frac{1}{1+\frac{[aTF]}{K_{D,1}}}$

An expression for $[aTF]$ is then needed to compute $[DNA]/[DNA]_T$

From (5):

$$[aTF] = [aTF]_T - [L \cdot aTF] - [DNA \cdot aTF]$$

Expressions for $[L \cdot aTF]$ and $[DNA \cdot aTF]$ are then required.

From (4):

$$[L \cdot aTF] = [L]_T - [L]$$

Using (1) to find an expression for $[L]$, the above can be re-written as:

$$[L \cdot aTF] = [L]_T - \frac{K_{D,L}[L \cdot aTF]}{[aTF]}$$

Combining terms in the above expression gives:

7) $[L \cdot aTF] = \frac{[L]_T}{\left(1+\frac{K_{D,L}}{[aTF]}\right)} = \frac{[L]_T[aTF]}{K_{D,L}+[aTF]}$

Similarly, using (3) and (2) gives:

$$[DNA \cdot aTF] = [DNA]_T - [DNA] = [DNA]_T - \frac{K_{D,1}[DNA \cdot aTF]}{[aTF]}$$

Therefore,

8) $[DNA \cdot aTF] = \frac{[DNA]_T[aTF]}{K_{D,1}+[aTF]}$

Using (7) and (8) in the expression for $[\mathrm{aTF}]$ gives:

$$0 = [\mathrm{aTF}] + \frac{[\mathrm{L}]_\mathrm{T}[\mathrm{aTF}]}{\mathrm{K}_{\mathrm{D,L}}+[\mathrm{aTF}]} + \frac{[\mathrm{DNA}]_\mathrm{T}[\mathrm{aTF}]}{\mathrm{K}_{\mathrm{D,1}}+[\mathrm{aTF}]} - [\mathrm{aTF}]_\mathrm{T}$$

This non-linear equation can be numerically solved for the single unknown $[\mathrm{aTF}]$, which can then be used in equation (6) to find f. The remaining unknowns can then be found from equations (1) through (5).

For studying the tetracycline-TetR system, we assume in this model that:

- TetR exists only as a dimer.
- Two tetracycline molecules bind to one TetR dimer.
- TetR dimer bound to tetracycline cannot bind DNA.
- One *tetO* site per DNA.
- The fraction of DNA unbound by TetR is proportional to the transcription output signal generated.

**SI Table 2: Parameters used for modeling the tetracycline-tetR system**

| Parameter | Value | Units | Reference and Note |
|---|---|---|---|
| $\mathrm{K}_{\mathrm{D,L}}$ | 7.94e-13 | M | [1] **a** |
| $\mathrm{K}_{\mathrm{D,1}}$ | 1.35e-11 | M | [2] **b** |
| $[\mathrm{DNA}]_\mathrm{T}$ | Variable | M | |
| $[\mathrm{L}]_\mathrm{T}$ | Variable | M | |
| $[\mathrm{aTF}]_\mathrm{T}$ | Variable | M | |

Notes:

a. Taken as the reciprocal of association constant for aTc and TetR, which is stated in [1].
b. Taken as the reciprocal of association constant for TetR and *tetO* site on DNA, which is computed per the correlation in [2] for the *tet* $O_2$ operator at 20 mM NaCl. This correlation is $\log \mathrm{K}_{\mathrm{as}} = 5.6 - 3.1 \log[\mathrm{Na}^+]$, where $[\mathrm{Na}^+] = 0.020\ \mathrm{M}$ in our experimental setup.

It is important to note that TetR is a homodimer, and that each homodimer has the capacity to repress one DNA template and has the capacity to bind two tetracycline molecules. Therefore, all expressions stated here relating to aTF concentration including $[\mathrm{aTF}]_\mathrm{T}$, $[\mathrm{aTF}]$, and $[\mathrm{L} \cdot \mathrm{aTF}]$ consider the TetR homodimer. When solving the generalized double equilibrium model stated in equations 1-8, the two-to-one ratio of tetracycline to TetR homodimer needs to be accounted for. To account for this, we use an “effective” total ligand concentration in the associated Jupyter Notebook calculations, which is equal to the total concentration of tetracycline added divided by two. When translating the model to other ligand-aTF systems, it is important to adjust the Jupyter Notebook calculations accordingly.

We then use this formalism to calculate fraction of unoccupied DNA template as a function of total DNA, TetR and aTc concentrations. A Jupyter Notebook solving this system of equations can be found in Supplementary_Data_2.ipynb

**Supplementary Files:**

**Supplementary_Data_1.xls**

DNA and protein sequences used in this study.

**Supplementary_Data_2.ipynb**

Jupyter notebook codes used to simulate results shown in Figure. 6.

**All_Source_Data.xls**

Source data for all figures of this study.
