## Supplementary material for "Engineering a cell-free biosensor signal amplification circuit with polymerase strand recycling": Jupyter Notebook Code for Supplementary Note: Supplementary_Data_2.ipynb

```
# %% [markdown]
## Supplementary Information for Engineering a cell-free biosensor signal amplification circuit with polymerase strand recycling

Yueyi Li$^{1,2}$, Tyler J. Lucci$^{1,2}$, Jaeyoung K. Jung$^{1,2,3}$, Matias V. Dujovne$^{4}$, Daiana A. Capdevila$^{4}$, Julius B. Lucks$^{1,2,3,5,*}$

1 – Department of Chemical and Biological Engineering, Northwestern University, Evanston, IL, USA.<br>
2 – Center for Synthetic Biology, Northwestern University, Evanston, IL, USA.<br> 
3 – Center for Water Research, Northwestern University, Evanston, IL, USA<br>
4 - Fundación Instituto Leloir, Instituto de Investigaciones Bioquímicas de Buenos Aires (IIBBACONICET), C1405BWE Ciudad Autónoma de, Buenos Aires, Argentina<br>
5 – Center for Engineering Sustainability and Resilience, Northwestern University, Evanston, Illinois 60208, USA<br> 
*Corresponding author: Julius B. Lucks<br> 

# %% [markdown]
##  Double equilibrium model to predict increasing sensitivity with PSR
The equation below is used to solve for [aTF] in the double equilibrium model for predicting dose-response of the ligand-aTF system. Remaining variables are then solved for via back substitution. **See Supplementary Information for the full model and equations** <br> 
$0 = [aTF] + \frac{{[L]_T}[aTF]}{K_{D,L}+[aTF]} + \frac{{[DNA]_T}[aTF]}{K_{D,1}+[aTF]} - {[aTF]_T}$ <br>

# %% [code]
import numpy as np
import pandas as pd
import matplotlib
import matplotlib.pyplot as plt
import matplotlib.font_manager as font_manager
from scipy.optimize import root_scalar
import seaborn as sns

# %% [code]
#returns x-value that makes function (y-values) 0 by linear interpolation
def interpolate_zero(x_vector,y_vector):
    val_zero = 0
    for k in range(1,len(x_vector)-1):
        y_ = y_vector[k]
        y_prev = y_vector[k-1]
        x_ = x_vector[k]
        x_prev = x_vector[k-1]
        #if function value is zero at this x, return the value of that x
        if y_ == 0:
            return x_
        #if function value is positive but previous value was negative, interpolate 
        elif y_ >= 0 and y_prev < 0:
            val_zero = x_prev+(0-y_prev)*(x_-x_prev)/(y_-y_prev)
            return val_zero
        #if function value is negative but previous value was positive, interpolate 
        elif y_ <= 0 and y_prev > 0:
            val_zero = x_prev+(0-y_prev)*(x_-x_prev)/(y_-y_prev)
            return val_zero
    return 'val_zero'

# %% [code]
#returns value for the double-equilibrium function to be zeroed 
def f_zero(x, bb, cc, dd, ee, ff):
    #[aTF] = x
    #[L]_T = bb
    #KDL = cc
    #[DNA]_T = dd
    #KD1 = ee
    #[aTF]_T = ff
    return x+bb*x/(cc+x)+dd*x/(ee+x) - ff

# %% [code]
#The below values for KDL and KD1 are specific to the tetracycline/tetR system
#If it is desired to study a different ligand/aTF system, simply change the KDL and KD1 values to those 
#corresponding to the desired ligand/aTF system

KDL = 1/(1.26e12)*10**9; #nM PMID 14764926

#KD1 is computed from the association constant using the correlation described in PMID 10071936
#This correlation is log⁡(K_as)=5.6-3.1 log⁡[Na+], where [Na+]=0.020 M in our experimental setup 
KD1 = 1/(10**(5.6 - 3.1*np.log10(20/1000)))*10**9 #nM PMID 10071936, 20 mM NaCl

print("KDL={:e} nM".format(KDL))
print("KD1={:e} nM".format(KD1))

#aTF_T: total aTF concentration. The total aTF concentration is the 
#total concentration of 'functional' aTF. In the case of aTFs that 
#dimerize this would be the concentration of aTF dimer used in the system.
aTF_T = 50 #nM, TMSD condition
#aTF_T = 25 #nM, PSR condition

#L_T: total ligand concentration
apoints = 31
L_T_vector = np.logspace(-2,5,apoints) #nM

#DNA_T: total DNA template concentration
DNA_T = 10; #nM, TMSD condition
#DNA_T = 1; #nM, PSR condition 

#convert L_T from nM to uM for plotting purposes
L_T_for_plot = []
for i in range(0,len(L_T_vector)):
    L_T_for_plot.append(L_T_vector[i]/1000)

fig, ax = plt.subplots(figsize = (8,5))
ax.patch.set_facecolor('white')
ax.patch.set_alpha(0)

#solve nonlinear equation for [aTF], backsolve for all other variables 
vals = []
for j in range(0,len(L_T_vector)):
    L_T = L_T_vector[j]
    L_T = L_T/2 #need two aTC per one TetR
    #In the case of TetR and SmtB dimers, two molecules of inducer are needed to de-activate the system.
    #We therefore divide total ligand concentration by two to get the 'effective' concentration of ligand
    #needed to de-activate the functional aTF unit.
        
    #compute aTF concentration via linear interpolation for use as an initial guess for Scipy root_scalar
    xvals = np.logspace(-21,np.log10(aTF_T)+1,1001)
    yvals = []
    for i in range(0,len(xvals)):
        aTF = xvals[i]
        function_value = f_zero(aTF, L_T, KDL, DNA_T, KD1, aTF_T)
        yvals.append(function_value)
    initial_guess = interpolate_zero(xvals,yvals)
    
    #compute aTF concentration via Scipy root_scalar using the initial guess generated via linear interpolation 
    sol = root_scalar(f_zero, args=(L_T, KDL, DNA_T, KD1, aTF_T), bracket=[0, aTF_T],
                     x0 = initial_guess)
    
    #compare which solution (root_ scalar or initial_guess) is better (i.e. closer to 0)
    aTF = initial_guess
    func_value_for_initial_guess = f_zero(initial_guess, L_T, KDL, DNA_T, KD1, aTF_T)
    func_value_for_root_scalar_root = f_zero(sol.root, L_T, KDL, DNA_T, KD1, aTF_T)
    if func_value_for_root_scalar_root**2 < func_value_for_initial_guess**2:
        aTF = sol.root
    
    #backsolve for all other species concentrations
    L = L_T/(1+aTF/KDL) #[ligand] = ...
    LaTF = L*aTF/KDL #[ligand*aTF] = ...
    DNA = DNA_T/(1+aTF/KD1) #[DNA] = ...
    DNAaTF = aTF*DNA/KD1 #[DNA*aTF] = ...
    vals.append(DNA/DNA_T) #[DNA]/[DNA_T] = normalized response

#plot dose-response curve for specific DNA concentration
label = f'Model\nDNA_T: {str(DNA_T)} nM,\n aTF_T: {str(aTF_T)} nM'
ax.set_xlabel('Ligand Concentration (\u03BCM)')
ax.set_ylabel('Normalized Response ($\mathregular{[DNA]/[DNA]_{T}}$)')
ax.plot(L_T_for_plot, vals, linewidth = 2, label = label, marker = 'o')
ax.set_xscale('log')
leg = ax.legend(frameon=False,loc='lower right')
plt.show()

#Save model output as csv (for Mac OS, this will save to the notebook's directory)
data_dict1 = {'[L] (µM)': L_T_for_plot, '[DNA]/[DNA]_T': vals}
df = pd.DataFrame(data_dict1)
df.to_csv('DE_Model1.csv')

# %% [code]
#The below values for KDL and KD1 are specific to the tetracycline/tetR system
#If it is desired to study a different ligand/aTF system, simply change the KDL and KD1 values to those 
#corresponding to the desired ligand/aTF system

KDL = 1/(1.26e12)*10**9; #nM PMID 14764926

#KD1 is computed from the association constant using the correlation described in PMID 10071936
#This correlation is log⁡(K_as)=5.6-3.1 log⁡[Na+], where [Na+]=0.020 M in our experimental setup 
KD1 = 1/(10**(5.6 - 3.1*np.log10(20/1000)))*10**9 #nM PMID 10071936, 20 mM NaCl

print("KDL={:e} nM".format(KDL))
print("KD1={:e} nM".format(KD1))

#aTF_T_vector: vector total aTF concentrations desired for study. The total aTF concentration is the 
#total concentration of 'functional' aTF. In the case of aTFs that 
#dimerize this would be the concentration of aTF dimer used in the system.
aTF_T_vector = [250,25,2.5,0.25,0.025] #nM

#L_T: total ligand concentration
apoints = 31
L_T_vector = np.logspace(-3,4,apoints) #nM

#DNA_T: ratio of total DNA template to aTF concentration
aTF_T_to_DNA_T_ratio = 25; #unitless

#convert L_T from nM to uM for plotting purposes
L_T_for_plot = []
for i in range(0,len(L_T_vector)):
    L_T_for_plot.append(L_T_vector[i]/1000)

fig, ax = plt.subplots(figsize = (8,5))
ax.patch.set_alpha(0)

data_dict2 = {'[L] (uM)': L_T_for_plot}
for i in range(0,len(aTF_T_vector)):
    aTF_T = aTF_T_vector[i]
    DNA_T = aTF_T/aTF_T_to_DNA_T_ratio
    #solve nonlinear equation for [aTF], backsolve for all other variables 
    vals = []
    for j in range(0,len(L_T_vector)):
        L_T = L_T_vector[j]
        L_T = L_T/2 #need two aTC per one TetR
        #In the case of TetR and SmtB dimers, two molecules of inducer are needed to de-activate the system.
        #We therefore divide total ligand concentration by two to get the 'effective' concentration of ligand
        #needed to de-activate the functional aTF unit.
        
        #compute aTF concentration via linear interpolation for use as an initial guess for Scipy root_scalar
        xvals = np.logspace(-21,np.log10(aTF_T)+1,1001)
        yvals = []
        for i in range(0,len(xvals)):
            aTF = xvals[i]
            function_value = f_zero(aTF, L_T, KDL, DNA_T, KD1, aTF_T)
            yvals.append(function_value)
        initial_guess = interpolate_zero(xvals,yvals)

        #compute aTF concentration via Scipy root_scalar using the initial guess generated via linear interpolation 
        sol = root_scalar(f_zero, args=(L_T, KDL, DNA_T, KD1, aTF_T), bracket=[0, aTF_T],
                         x0 = initial_guess)

        #compare which solution (root_ scalar or initial_guess) is better (i.e. closer to 0)
        aTF = initial_guess
        func_value_for_initial_guess = f_zero(initial_guess, L_T, KDL, DNA_T, KD1, aTF_T)
        func_value_for_root_scalar_root = f_zero(sol.root, L_T, KDL, DNA_T, KD1, aTF_T)
        if func_value_for_root_scalar_root**2 < func_value_for_initial_guess**2:
            aTF = sol.root

        #backsolve for all other species concentrations
        L = L_T/(1+aTF/KDL) #[ligand] = ...
        LaTF = L*aTF/KDL #[ligand*aTF] = ...
        DNA = DNA_T/(1+aTF/KD1) #[DNA] = ...
        DNAaTF = aTF*DNA/KD1 #[DNA*aTF] = ...
        vals.append(DNA/DNA_T) #[DNA]/[DNA_T] = normalized response

    #plot dose-response curve for specific DNA concentration
    label = f'DNA_T: {str(DNA_T)} nM,\n aTF_T: {str(aTF_T)} nM'
    ax.plot(L_T_for_plot, vals, linewidth = 2, label = label, marker = 'o')
    data_dict2[label] = vals
    

ax.set_xlabel('Ligand Concentration (\u03BCM)')
ax.set_ylabel('Normalized Response ($\mathregular{[DNA]/[DNA]_{T}}$)')
ax.set_xscale('log')
leg = ax.legend(frameon=False,loc='center left', bbox_to_anchor=(1, 0.5))
plt.show()

#Save model output as csv (for Mac OS, this will save to the notebook's directory)
df = pd.DataFrame(data_dict2)
df.to_csv('DE_Model2.csv')
    

# %% [code]
```
